# Assessment of the World Largest Afforestation Program: Success, Failure, and Future Directions

**DOI:** 10.1101/105619

**Authors:** J. J. Zhu, X. Zheng, G.G. Wang, B.F. Wu, S.R. Liu, C.Z. Yan, Y. Li, Y.R. Sun, Q.L. Yan, Y. Zeng, S.L. Lu, X. F. Li, L.N. Song, Z. B. Hu, K. Yang, N.N. Yan, X.S. Li, T. Gao, J. X. Zhang, Aaron M. Ellison

## Abstract

The Three-North Afforestation Program (TNAP), initiated in 1978 and scheduled to be completed in 2050, is the world’s largest afforestation project and covers 4.07 x 10^6^ km^2^ (42.4%) of China. We systematically assessed goals and outcomes of the first 30 years of the TNAP using high-resolution remote sensing and ground survey data. With almost 23 billion dollars invested between 1978 and 2008, the forested area within the TNAP region increased by 1.20 × 10^7^ ha, but the proportion of high quality forests declined by 15.8%. The establishment of shelterbelts improved crop yield by 1.7%, much lower than the 5.9% expected once all crop fields are fully protected by shelterbelts. The area subjected to soil erosion by water decreased by 36.0% from 6.72 × 10^7^ to 4.27 × 10^7^ ha; the largest reductions occurred in areas where soil erosion had been most severe and forests contributed more than half of this improvement. Desertification area increased from 1978 to 2000 but decreased from 2000 to 2008; the 30-year net reduction was 13.0% (4.05×10^6^ ha), with 8.0% being accounted for by afforestation in areas with only slight, prior desertification. In addition to its direct impacts, the TNAP has enhanced people’s awareness of environmental protection and attracted consistent attention and long-term commitment from the Chinese government to the restoration and protection of fragile ecosystems in the vast Three-North region. The significant decline in forest quality, limited success in reducing desertification, and low coverage of shelterbelts are aspects of the TNAP in need of re-assessment, and additional ca. 34 billion dollars will be needed to ensure the completion of the TNAP.

## Introduction

Large-scale afforestation programs have been attempted worldwide to improve environmental conditions in deteriorated or unfavorable sites (Zhu, 2013). Most of the largest attempts did not conclude successfully for social-political, technical, or ecological reasons; examples include (Stalin’s) Great Plan for the Transformation of Nature in the former Soviet Union (Brain, 2010); the Great Plains Shelterbelt Project in the USA (Orth, 2007); and the Green Dam Engineering Project in five North African countries (Jiang et al., 2003). However, the world’s most ambitious afforestation program, China’s Three-North Afforestation Program (TNAP), is still ongoing.

The TNAP (also known as the Three North Forest Protection System-China [Moore and Russell, 1990] and the Three-North Protective Forest Program [Fang et al., 2001; Zhu et al., 2009]) was initiated in 1978 and is scheduled to be completed by 2050. This afforestation plan covers ≈4.07 × 10^6^ km^2^ (> 42%) of the land area of China, and encompasses almost all of the country’s arid and semiarid land areas (Fig. 1A). The central government of China invested 23 billion dollars into the TNAP between 1979 and 2008 (Table AS1), in the hopes that the construction of this “Great Green Wall” would greatly improve the environment in China’s “Three-North” regions (i.e., Northeast, Northwest, and North Central). Specific ecological benefits of increased forest cover were anticipated to include protection of agriculture and animal husbandry; reduction of soil erosion; and control of desertification. These enhanced ecological benefits are expected, in turn, to contribute to reductions in poverty, improvements in livelihoods, and positive restructuring of rural economies (Liu et al., 2009).

**Fig. 1.**
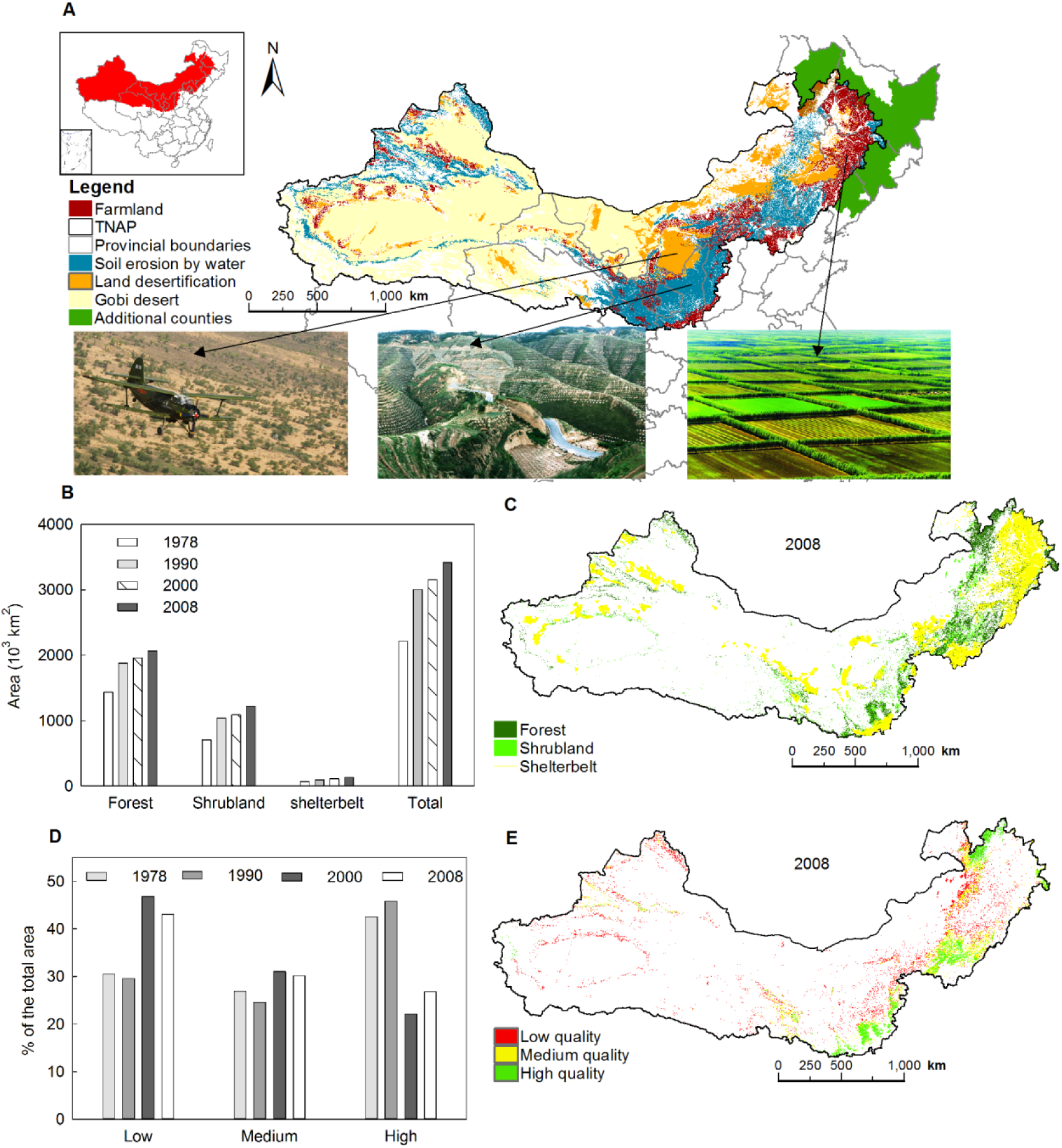
(A) The spatial distribution of farmland, desertification, and soil erosion areas, with photographs from the China Forestry News (used with permission) illustrating the corresponding forest types in 1978, with an insert map showing the TNAP area subjected to this study (in red) and areas added in 2000 (in green); (B) the change of forest area from 1978 to 2008; (C) the spatial distribution of different forest types in 2008; (D) change in the proportion of forests of different quality types from 1978 to 2008; (E) the spatial distribution of different forest quality types in 2008.

Despite the consistent support from the Chinese central government, the rationale and success of the TNAP have been questioned by scientists and non-specialists alike. On the one hand, several studies suggested that planting trees in arid and semi-arid regions, where annual precipitation < 400 mm, has exacerbated environmental degradation (Cao, 2008, Cao, 2011; Cao et al., 2010a, b). Sun et al. (2006) also reported that the water yields had dropped by 30–50% and vegetation cover had decreased by 6% after the TNAP was implemented in the Loess Plateau region (Fig. S1). On the other hand, a recent assessment in the Horqin Sandy Land (a semi-arid sandy region and the more representative area in terms of afforestation in TNAP region, Fig. S1) found that the TNAP played a significant role in controlling desertification expansion in the study area (Yan et al., 2011).

More than half way through its implementation, however, no one has systematically evaluated the outcomes of the TNAP in light of its three central goals: 1) increasing crop yield by protecting farmland with shelterbelts; 2) reducing soil erosion; and 3) controlling desertification. In this study, we systematically evaluated how the TNAP changed forest quantity and quality during its first 30 years, and how these changes impacted crop yield, soil erosion and desertification. By identifying successes and revealing failures, our assessment illustrates how to refine and improve the next 34 years of the TNAP.

### Forested quantity and quality

We estimated the forested area within the TNAP region using both remote sensing (Landsat MSS/TM/ETM+ and SPOT5) and field survey data. Our estimates from remotely-sensed data were 86.0% accurate based on validation with field data. Total forested area, including forests (canopy cover ≥30%, minimum area > 400 m^2^), shrublands (canopy cover ≥40%), and shelterbelts (length >20 m), increased steadily from 220,995 km^2^ (5.5% of total land cover) in 1978, to 300,265 km^2^ (7.5%) in 1990, 315,280 km^2^ (7.9%) in 2000, and 341,391 km^2^ (8. 6%) in 2008. By 2008, there were 13,031 km^2^ of shelterbelts around farmlands; 178,985 km^2^ of forests and 89,749 km^2^ of shrublands in areas prone to soil erosion; and 27,402 km^2^ of forests and 32,224 km^2^ of shrublands in areas subject to desertification (Fig. 1A, B and C).

We derived percent canopy cover from NOAA AVHRR Normalized Difference Vegetation Index (NDVI) and MODIS NDVI, and used these values to classify forests and shrublands into three quality classes: high (canopy cover: > 55.0%), medium (forest canopy cover: 40.0-55.0% or shrubland canopy cover: 45.0-55.0%), and low (forest canopy cover: 30.0-40.0% or shrubland canopy cover: 40.0-45.0%). The proportion of the high-quality forests and shrublands decreased from 43% in 1978 to 27% in 2008, while the proportion of medium- and low-quality forests and shrublands increased, respectively, from 27% and 31% in 1978 to 30% and 43% in 2008 (Fig. 1D **and** E).

### Impact on crop yield

Shelterbelts or windbreaks are a critical component of the TNAP and have been established to improve microclimatic conditions for crop growth (Cao, 1983; Zhu, 2008). We first divided the farmland in the TNAP region into high, medium, and low climatic potential productivity zones based on solar radiation, temperature and precipitation (Fig. 2A). Within each zone, we then determined the relationships between maize yield and the level of protection from shelterbelts (Zheng et al., 2016) (Fig. 2B), and calculated the average level of protection by shelterbelts (Fig. 2C). Based on the regression equation developed for each zone (Fig. 2B), we estimated the maize yield under different levels of protection from shelterbelts.

**Fig. 2.**
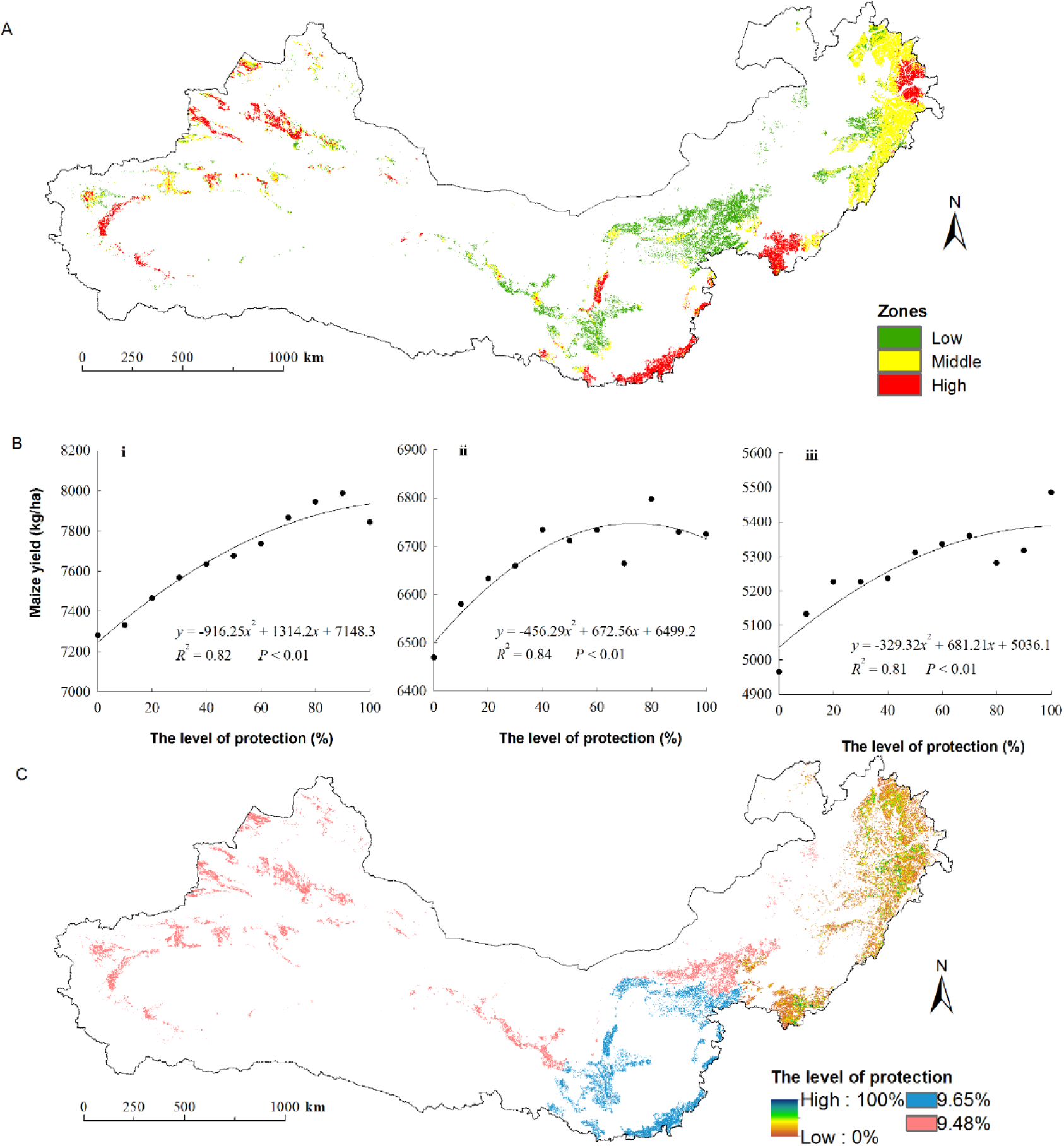
The distribution of low-, medium-, and high- potential productivity zones (A), the relationship between maize yield and the level of shelterbelt protection of farmland (B; i = high-, ii = medium-, and iii = low- potential productivity zones), and the spatial distributions of the level of protection of farmland in the TNAP region (C).

Maize yield increased steadily with the increasing level of shelterbelt protection, asymptoting at ≈80% (Fig. 2B). With the protection level at 80%, we calculated that shelterbelts could potentially improve the crop yield by nearly 6.0% (5.3%, 4.4% and 8.0% in the high-, medium-, and low-climatic potential productivity zones, respectively). However, the actual level of shelterbelt protection for farmland in the TNAP region averaged only 14.0% (13.3%, 17.7%, and 10.9% for the high-, medium-, and low-climatic potential productivity zones, respectively). After 30 years, therefore, shelterbelts in the TNAP region have improved crop yields by < 2.0% (2.2%, 1.6% and 1.4% for the high-, medium-, and low-climatic potential productivity zones, respectively).

### Impact on soil erosion

Soil erosion is a widespread problem throughout China, especially in the Loess Plateau region (Fig. S1). To assess the effects of the TNAP on soil erosion, we first estimated the area and intensity of soil erosion in the TNAP region using the revised universal soil loss equation (RUSLE), and estimated its parameters from local rainfall, topography, soil classification, and remote sensing data. The estimated annual soil erosion area in 2008 matched well with the Bulletin of Soil and Water Loss in China in 2008 (The Ministry of Water Resources of the People’s Republic of China, 2009). The estimated intensity of soil erosion in 2008 also agreed well with the monitored values from 20 hydrological observation stations (*R*
^
*2*
^ = 0.73, RMSE = 1485 t ha^-1^ year^-1^). To determine the variables affecting the change of soil erosion, we first divided the TNAP region into 5057 25 km × 25 km cells for path analysis. We then calculated the area and intensity of soil erosion, evapotranspiration (1-km resolution), the areas of forests and shrubland (30-m resolution), and the precipitation for each cell (25-km resolution). Finally, we used path analysis to determine the contribution of forests and shrublands to the reduction of soil erosion.

After 30 years of the TNAP, the areas of very slight, slight, moderate, severe, and extremely severe soil erosion by water decreased by 24%, 34%, 50%, 83% and 99%, respectively (Fig. 3A **and** 3B). Forests and shrublands played an increasingly important role in reducing both the intensity and the area of soil erosion by water (Fig. 3C **and** 3D). The contribution of forests and shrublands to the reduction of soil erosion area was 53% (42% from forests and 11% from shrublands) during 1978-2008, 37% during 1978-1990, 40% during 1990-2000, and 39% during 2000-2008 (Fig. 3C, 3D).

**Fig. 3.**
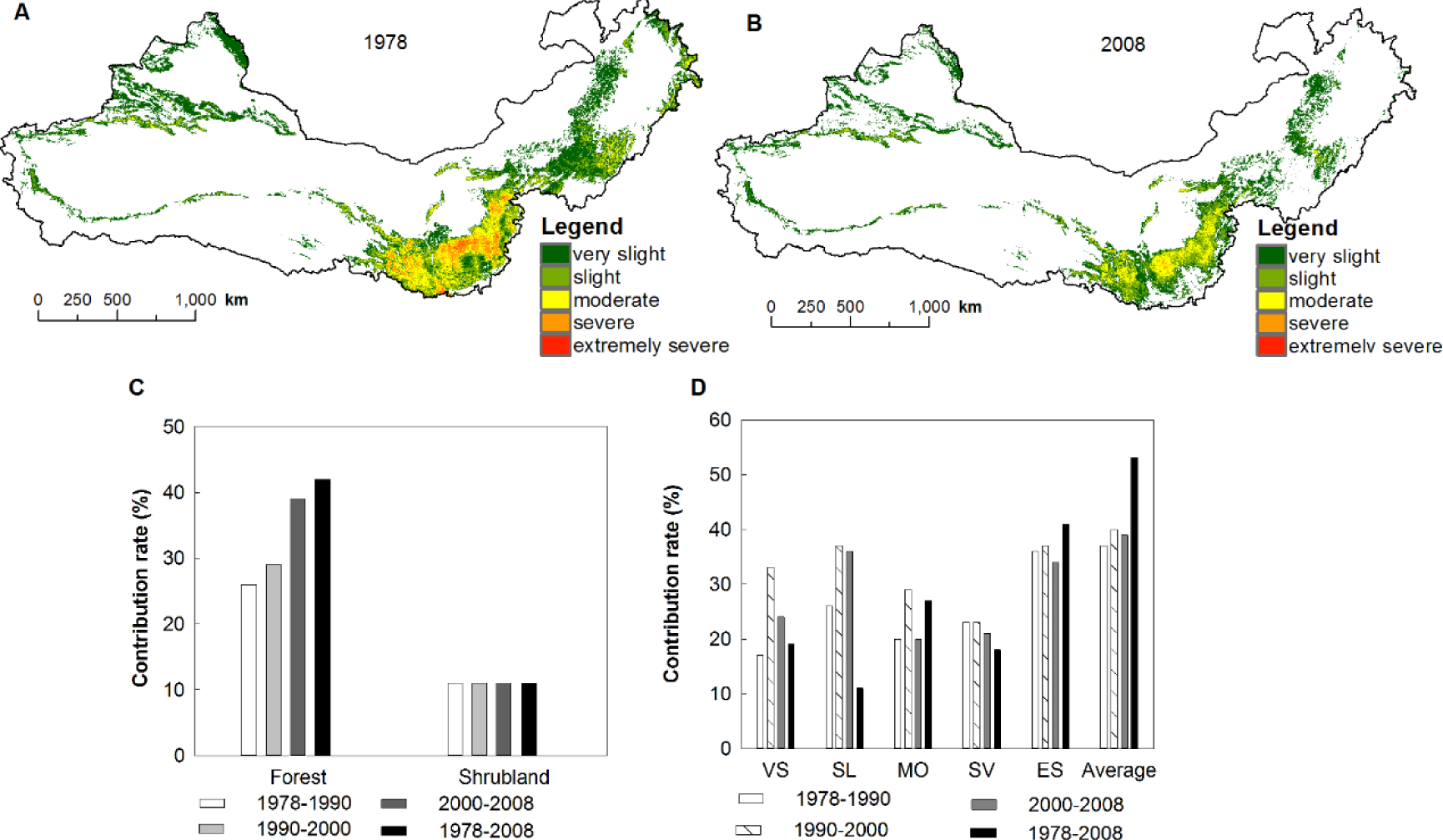
Soil erosion by severity class in 1978 (A) and 2008 (B) (the blank areas are the land without water-driven soil erosion); the contribution of forests and shrublands to the reduction of the total area of soil erosion (C); and to the different intensities of soil erosion (D) (VS: very slight, SL: slight, MO: moderate, SV: severe, ES: extremely severe soil erosion) during 1978-2008.

### Impact on desertification

Desertification in arid, semi-arid, and dry sub-humid regions is of global concerns because of its political and socio-economic ramifications (Reynolds et al. 2007) and 90% of China’s lands subject to aeolian desertification are within the TNAP region. We used the method of Yan et al. (2009) to classify areas in the TNAP region that are prone to aeolian desertification into four intensity levels (slight, moderate, severe, and extremely severe), and estimated the area of each using Landsat MSS, ETM+, and TM data from 1978, 1990, 2000 and 2008. Our estimates were 95% accurate based on field surveys (3,100 30 m×30 m plots). We defined the reduction of desertification area due to the TNAP based solely on site occupancy by forests and shrublands.

The area of the desertification increased from 310,589 km^2^ in 1978 to 343,108 km^2^ in 1990 and 375,609 km^2^ in 2000, and then decreased to 270,096 km^2^ in 2008 (Fig. 4A). However, the area of forests and shrublands on land subject to aeolian desertification increased steadily from 33,394 km^2^ in 1978 to 51,541 km^2^ in 1990, 57,267 km^2^ in 2000, and to 9,626 km^2^ in 2008. Over the 30 years, land subject to desertification declined by 40,494 km^2^ (13%) while forested area increased by 26,232 km^2^ (79%). The increase of forested area directly accounted for 65% of the net reduction of land area subject to desertification, but > 90% of the forests and shrublands had been successfully established only in areas classified as slight desertification (Fig. 4B). Considering the overall size of land subject to desertification, the establishment of forests and shrublands by the TNAP played at best a limited role in reducing desertification between 1978 and 2008 (Fig. 4C **and** D).

**Fig. 4.**
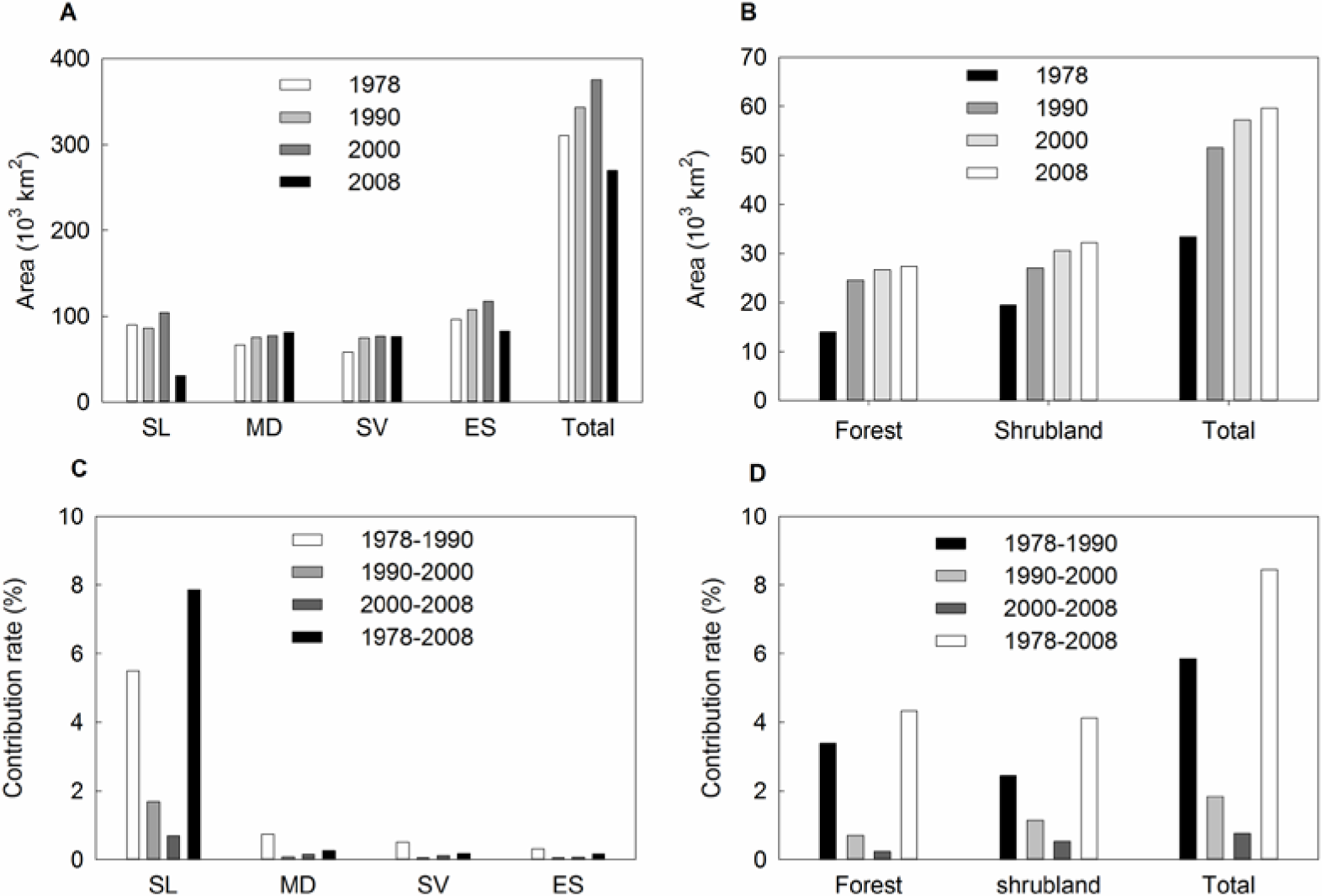
Changes in the areas of land subject to desertification in the TNAP region (A); changes in the areas of forests and shrublands in these lands (B); the contribution of forests and shrublands to the reduction of the four intensity levels of the desertification (C); and the contribution of forests and shrublands to the reduction of the total area of the desertification at all levels (D) during 1978-2008 (SL: slight, MO: moderate, SV: severe, ES: extremely severe desertification areas).

## Discussion

Forest cover in the TNAP region increased steadily from 5.5% in 1978 to 8. 6% in 2008. This increase can be attributed to both direct afforestation (e.g., planting trees) and natural recovery of forests following the implementation and enforcement of strict protection measures. However, the central government of China reported a much larger increase (5.1% in 1978 to 10.6% in 2008: Liu et al., 2009). Although part of the government’s overestimate is attributable to an underestimate of initial forest coverage, inconsistent definitions of forest cover are much more important. In 1978, forested areas were defined as those with canopy cover ≥30% for forests and ≥40% for shrublands (the same definitions we use here), but since 1994, the government has defined forested areas as those with canopy cover ≥20% for forests and ≥30% for shrublands, which together inflate current estimates of overall forest cover. After revising the government’s data to reflect a more accurate initial value and by using a consistent (and more stringent) definition of forest cover, their estimates differ from ours by < 1%. Although our estimates of forest cover use identical boundaries for the TNAP region (as defined in 1978: 551 counties across 13 provinces; Fig.1 inset map in red), the government report expanded the boundary of the TNAP region to include 600 counties across the 13 provinces in 2000 (Fig. 1 inset map in green). This 10% expansion also could have contributed to higher forest cover estimates in the government report because the 49 additional counties included since 2000 have climatic and site conditions more suitable for forest growth.

Both our estimate and the government’s estimate show an increase in forest cover, but forest quality declined from 1978 to 2008 (Fig. 1D, E). The higher initial forest quality likely resulted from the preponderance of natural forests in the TNAP region in 1978. As forests planted during the TNAP were expanded into poor sites, especially when tree species were mismatched with planting site conditions (e.g., popular was planted in the arid area), low survival rate and poor growth of planted trees led to the development of the low-quality forests. If improving forest and shrubland quality is an important goal of the TNAP, better silvicultural practices for improving the quality of current forests and shrublands will be needed.

Shelterbelts planted during the TNAP increased crop yield by < 2%, far from the hoped-for increase of 6% (Fig. 2). The poor success of shelterbelts in meeting the productivity goal of the TNAP is most likely attributable to the low level of protection achieved (≈14%). Perhaps further gains in crop production could accrue if additional shelterbelts are planted in the next three decades. However, planting of TNAP shelterbelts largely occurred between 1978 and 1990, when farmland belonged to the government (Zheng et al., 2013). After 1990, the government adopted a household contract responsibility system, and leased farmland to the farmers. Even though shelterbelts can lead to increasing the overall production of crops at large scales (Cao, 1983; Jiang et al., 2003; Orth, 2007), small-scale farmers have been reluctant to plant new shelterbelts because they reduce the amount of available arable land and increases competition for nutrients, water, and light between trees and adjacent crops (Zheng et al., 2016). To resolve this contradiction, we suggest that local governments consider removing area designated for shelterbelts from farmland leases. Land used for irrigation ditches, roads, and other agricultural infrastructure also could be included in areas designated for shelterbelts (Cao, 1983). Finally, economic compensation should be provided to farmers whose farmland is located near the shelterbelts.

Forests and shrublands planted during the TNAP effectively reduced soil erosion (Fig. 3). More importantly, the reduction in soil erosion was most effective on the severe and extremely severe erosion classes. This very positive impact of the TNAP appears to have resulted from planting trees in areas with relatively high precipitation that both erodes exposed soils and improves plant growth. As the planted forests have developed, the rainfall intercepted by the canopy and taken up by the well-developed root systems have protected effectively the soil from erosion by water. However, these forests are still of relatively poor quality compared to those in similar precipitation zones elsewhere in the world. (Zheng and Zhu, 2013). In the next phase of the TNAP, improving forest quality through innovative silviculture could further enhance the role of this program in reducing soil erosion (Cao et al., 2011).

The contribution of forests and shrublands to the reduction of areas subject to aeolian desertification was small, and most of this reduction occurred in the areas classified as slight desertification (Fig. 4). Most of the areas at risk of aeolian desertification areas have too little rainfall to support forests or shrublands (Cao et al., 2010a, b) and it is unrealistic to expect the TNAP to reduce the severe desertification. Indeed, despite a steady increase of forested area from 1978 to 2008, areas subject to desertification actually increased from 1978 to 2000, and only decreased between 2000 and 2008, suggesting that factors other than planted trees were driving the expansion and contraction of desertification areas. In fact, the increase in desertification areas between 1978 and 2000 most likely was caused by estrepement, including over-grazing by livestock, denudation of land, and overuse of available groundwater resources (Cao, 2011).

In the areas of the TNAP region at risk of desertification, many established forests are declining because tree species selected for afforestation were poorly matched to local site conditions (Cao, 2008; Yan et al., 2011, Zhu, 2013) (Tab. AS2). Since water is the key factor limiting successful afforestation in these areas (Yan et al., 2011), water availability must be considered explicitly in the TNAP. We classify the TNAP region into four afforestation zones (Fig. 5) based on the aridity index (AI) developed by Zheng and Zhu (2016), and then select tree species for afforestation to match each zone (Tab. 1).

**Fig. 5.**
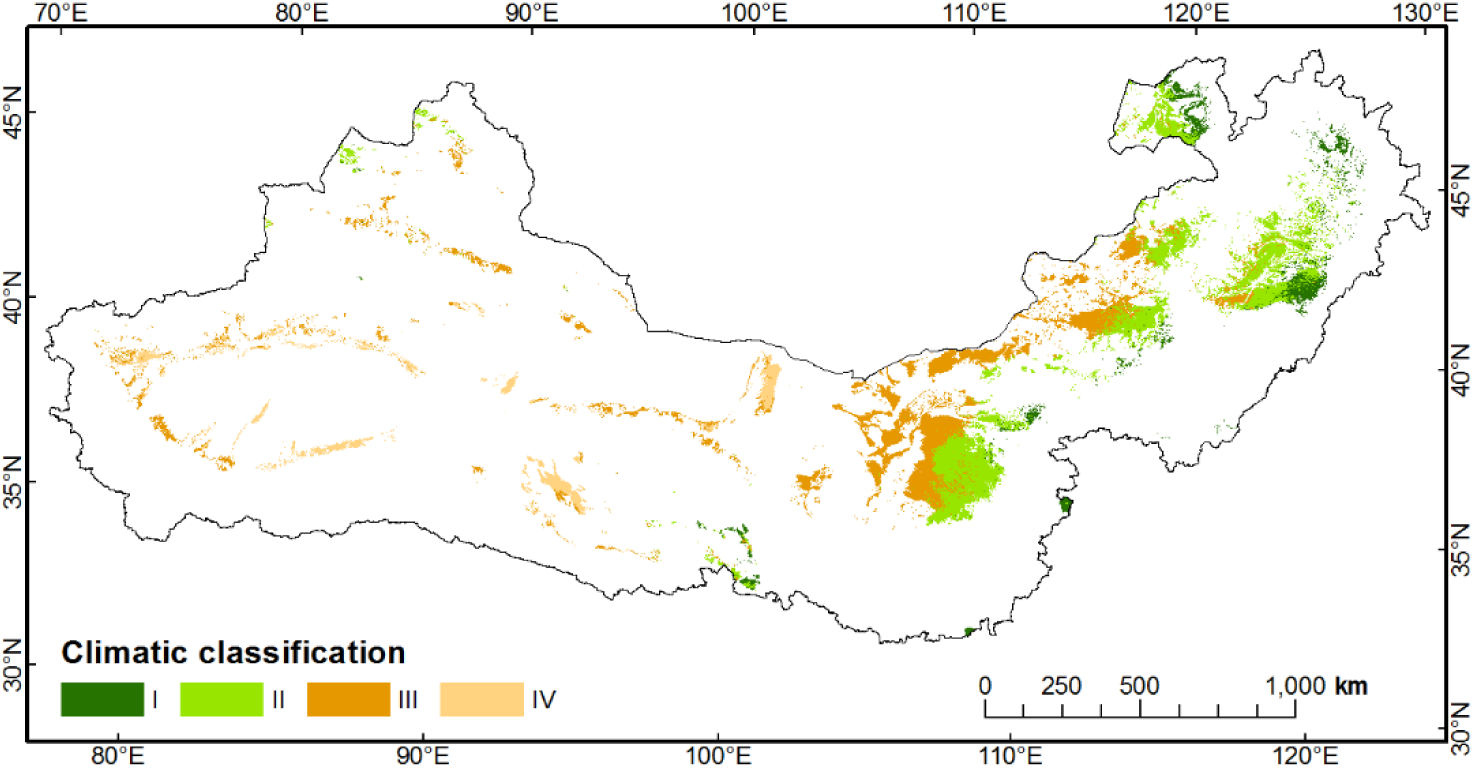
Climatic classification of afforestation in areas at risk of desertification in the TNAP region: semiarid and sub-humid-arbor zone (I); semiarid-shrub zone (II); arid-shrub steppe zone (III); hyperarid-no vegetation zone (IV).

**Table 1.**
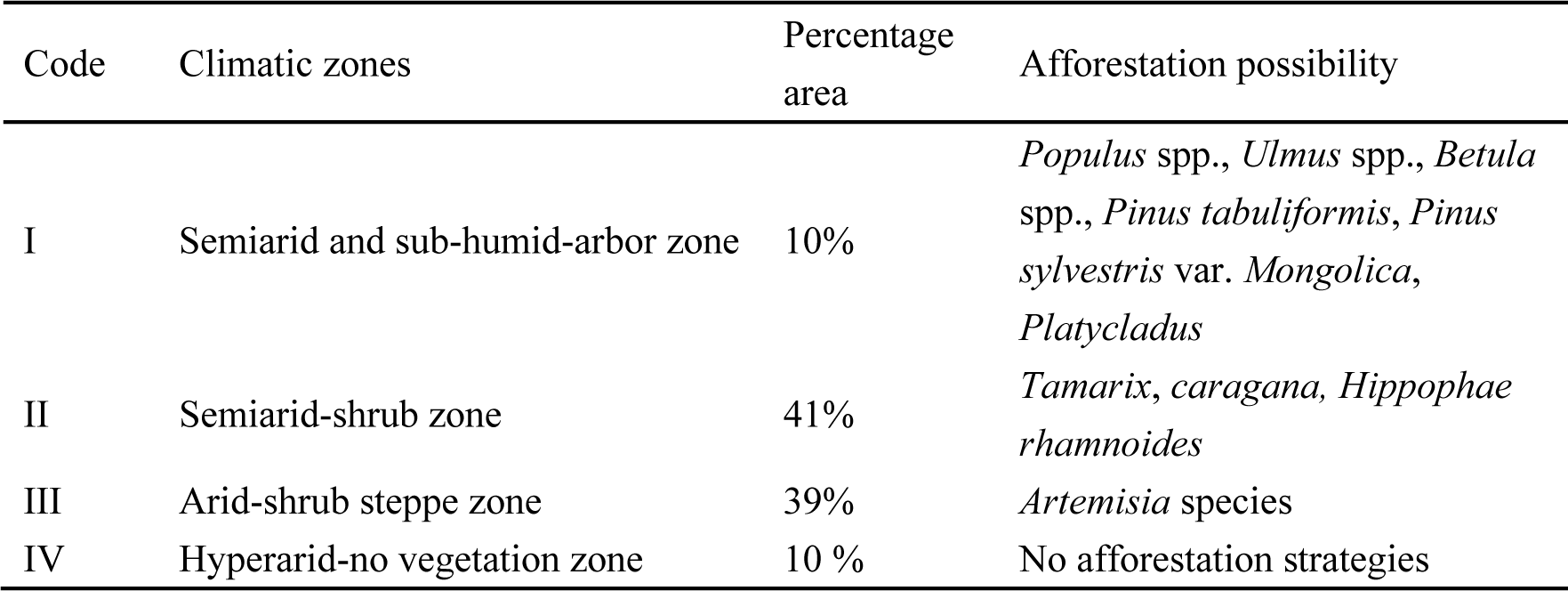
Descriptions of climatic zones and afforestation possibilities.

| Code | Climatic zones | Percentage area | Afforestation possibility |
| --- | --- | --- | --- |
| I | Semiarid and sub-humid-arbor zone | 10% | *Populus* spp., *Ulmus* spp., *Betula* spp., *Pinus tabuliformis*, *Pinus sylvestris* var. *Mongolica*, *Platycladus* |
| II | Semiarid-shrub zone | 41% | *Tamarix*, *caragana*, *Hippophae rhamnoides* |
| III | Arid-shrub steppe zone | 39% | *Artemisia* species |
| IV | Hyperarid-no vegetation zone | 10 % | No afforestation strategies |

We also note that our analysis does not differentiate between direct effect of the other government programs to reduce estrepement, or implementation and enforcement of environmental protection (Tab. AS3). An important positive outcome of the TNAP has been the increasing recognition by the Chinese central government of the importance of environmental protection. During the same time that the TNAP has been running, the government has implemented many other environmental programs, including the Natural Forest Conservation Project, the Returning Cropland to Forestry and Grass, the Sand Prevention Engineering Surrounding Beijing and Tianjin, the Wildlife Protection and Nature Reserve Development Program, as well as shelter forest projects around the country (including the Coastal Shelter Forest Project, the Comprehensive Management of Shelter Forest Project in the Pearl River Basin, Shelter Forest Project in the upper reaches of the Yangtze River). Together with the TNAP, these programs could potentially alleviate the pressure of rapid economic growth on the environment and help improve the future ecological security of China.

## Conclusions

Our range-wide assessment reveals that the TNAP has provided measurable benefits to the Three North regions. Forest cover increased, shelterbelts improved crop yields, and forests and shrublands reduced the amount and area of soil erosion. In addition to these measureable benefits, the TNAP likely had a profound influence on the policy and practice of environmental protection in China. Our assessment also highlights challenges for the remaining three decades of the TNAP. First, the quality of forests established by the TNAP must be improved. Second, land-tenure policies associated with small farms need to be changed to promote planting of additional shelterbelts. Third, controlling desertification by afforestation will be effective only in areas already at low risk of aeolian desertification, and species selection must be explicitly guided by site water availability. Finally, the first 30 years of TNAP has already cost 23 billion dollars (Tab. AS1), it is likely that an investment of one billion dollars will be needed each year to ensure the completion of the TNAP.

## Acknowledgments

We thank JY Fang, FL He, FQ Jiang and XG Han for their comments and suggestions on this manuscript. This work was supported by grants from CAS Key Research Program of Frontier Sciences (QYZDJ-SSW-DQC027), National Nature Science Foundation of China (31025007) and the Knowledge Innovation Program of Chinese Academy of Sciences (KZCX1-YW-08-02). AME’s participation was supported by the Chinese Academy of Sciences (CAS) Presidential International Fellowship Initiative for Visiting Scientists (2016VBA074).

## 4 Supplementary Materials

*Data collection*

*Definitions of forests and shrublands*

*Monitoring of afforestation area for forests, shrublands, and shelterbelts*

*Monitoring of the quality of forests, shrublands, and shelterbelts*

*Effects of shelterbelt on crop yields*

*Effects of forests and shrublands on soil erosion by water*

*Effects of forests and shrublands on desertification*

## Supplemental Results, Tables and Figures

*Afforestation areas/mapping of forests and shrublands from 1978 to 2008*

*The quality of forests and shrublands from 1978 to 2008*

*Impact on crop yields*

*Impact on soil erosion by water*

*Impact on desertification*

## Uncertainty analysis of the major assessment results

*Afforestation areas and the quality of forests, shrublands and shelterbelts*

*Impact on crop yields*

*Impact on soil erosion by water*

*Impact on desertification*

## References

### Additional tables

Table AS1 other government programs implemented in the Three North regions after the TNAP

Table AS2 other government programs implemented in the Three North regions after the TNAP

Table AS3 other government programs for vegetation restoration, which had or have been implemented in the Three North regions after the TNAP

## Materials and methods

### Study area

The Three-North region (TNR) of China (Northeast China, North China, and Northwest China) has a total area of 4.069 × 10^6^ km^2^. It is located between meridians 73°26′ E and 127°50′ E and parallels 33°30′ N and 50°12′ N, includes 551 counties across 13 provinces, and accounts for more than 42.4% of China’s total territory (1). The TNR is commonly divided into four zones: Northeast China, North China, Loess Plateau and Northwest China, according to geo-morphological and climatic characteristics (Fig. S1) (2, 3).

**Fig. S1.**
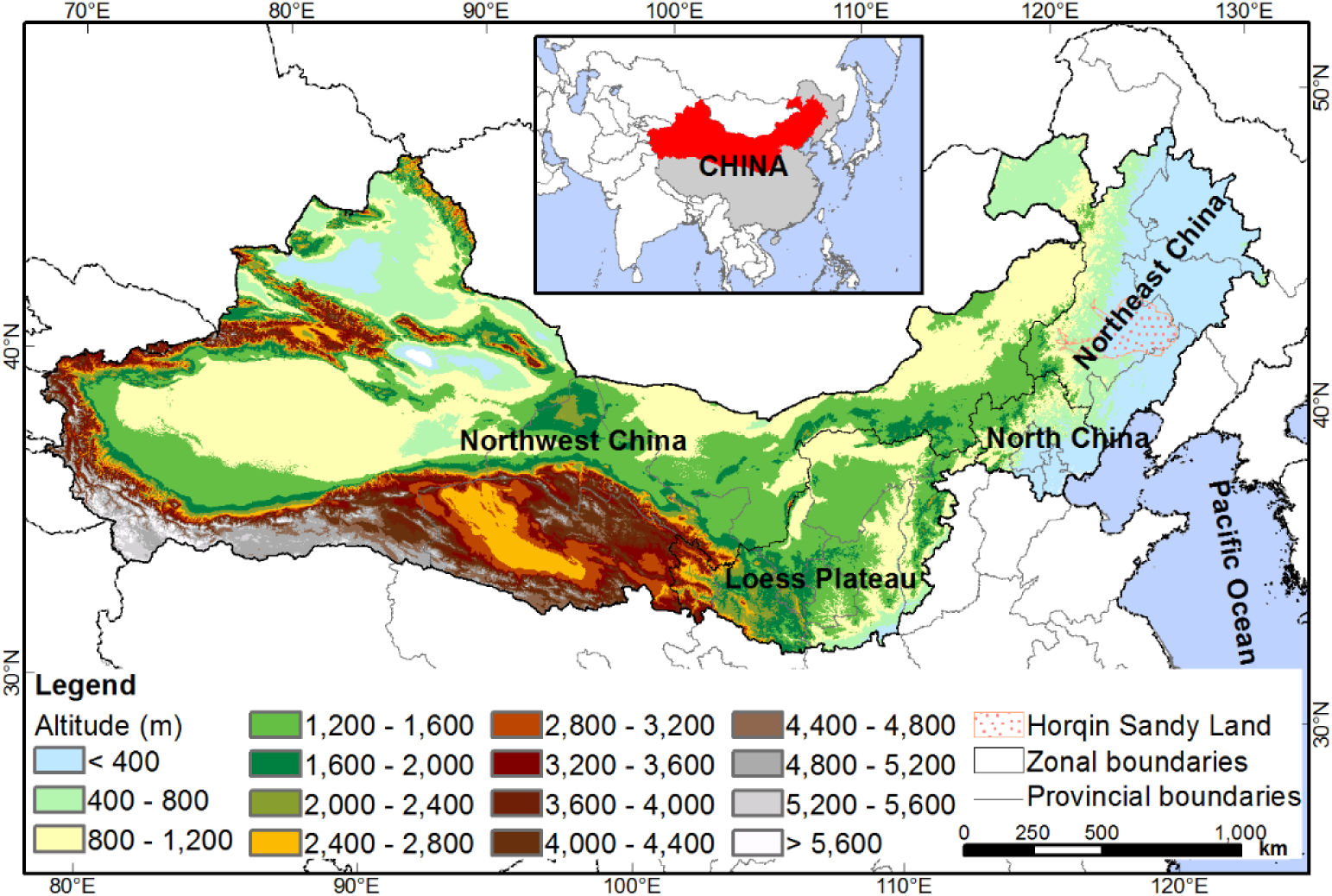
Study area location and the four zones of the Three-North regions.

### Definition of forested area

The forested areas in the Three-North Forestation Program (TNAP) include forests, shrublands, and shelterbelts or windbreaks. Areas with the tree canopy cover > 30% are defined as forests; areas with the shrub cover >40% are defined as shrublands; a single or multiple row of planted trees with the row length <u>> 20 m are defined as</u> shelterbelts (4).

### Datasets

#### Remote sensing images

Landsat series images: four time series (1978, 1990, 2000 and 2008) of high quality Landsat satellite images covering the Three-North regions (TNR) were collected during the growing seasons. First, 165 scenes from 1977-1980 were obtained from Landsat MSS images with a spatial resolution of 80 m to represent the year 1978. Second, 278 scenes from 1989 to 1992, 268 scenes from 1999 to 2001 and 258 scenes from 2007 to 2009 were obtained from Landsat TM with a spatial resolution of 30 m to represent the year 1990, 2000, and 2008, respectively. These images were collected from the USGS Landsat archive (http://glovis.usgs.gov) and the China remote sensing satellite ground station (www.ceode.cas.cn).

SPOT5: nine scenes of SPOT5 with a spatial resolution of 2.5 m were collected between 2007 and 2008 during the growing seasons, and these images were used to interpret the major forest types covering the TNR in 2008. These images were bought from the Beijing Shibao Satellite Image Co., Ltd. (http://www.spotimage.com.cn).

CBERS-02B HR: six scenes of CBERS-02B HR images with a spatial resolution of 2.36 m were collected from 2007 and 2009 to estimate the shelterbelt areas in 2008. These images were obtained from China Centre for Resources Satellite Data and Application (http://www.cresda.com/EN/).

TRMM: the Tropical Rainfall Measuring Mission (TRMM) is a joint mission between the National Aeronautics and Space Administration (NASA) and the Japan Aerospace Exploration Agency designed to monitor and study tropical rainfall. Version 7 TRMM images with a spatial resolution of 25 km covering the TNR from 2000 to 2009 were collected. TRMM 3B43 data were downloaded from the NASA Goddard Earth Sciences Data and Information Services Center website (http://mirador.gsfc.nasa.gov/).

MODIS: Moderate Resolution Imaging Spectroradiometer (MODIS) data were obtained from NASA MODIS sensors aboard the Terra satellite and were downloaded from the EOS Gateway at https://lpdaac.usgs.gov/. Annual composite MODIS GPP data (MOD17A3) at a 1-km spatial resolution from 2007 to 2009 were downloaded. Besides, MOD13A3 data between 2000 and 2008 containing the monthly normalized difference vegetation index (NDVI) were downloaded and used in this study.

#### Other data

Meteorological station data: observations of air temperature were obtained from ground meteorological stations. They were measured according to the standards of the World Meteorological Organization at 2 m above the ground. Temperature observations from 2001 to 2009 were used to estimate annual temperature and evapotranspiration. Annual precipitation observations from 2001 to 2009 were obtained from ground meteorological stations and were used to verify the accuracy of the estimated precipitation in a raster format with 1-km resolution grids.

Maize sample plot survey: fifty-eight experimental farmland plots with 1000 m × 1000 m area each were collected in the typical districts such as the counties of Dehui, Nong'an and Changling in Northeast China (Fig. S2) (5).

**Fig. S2.**
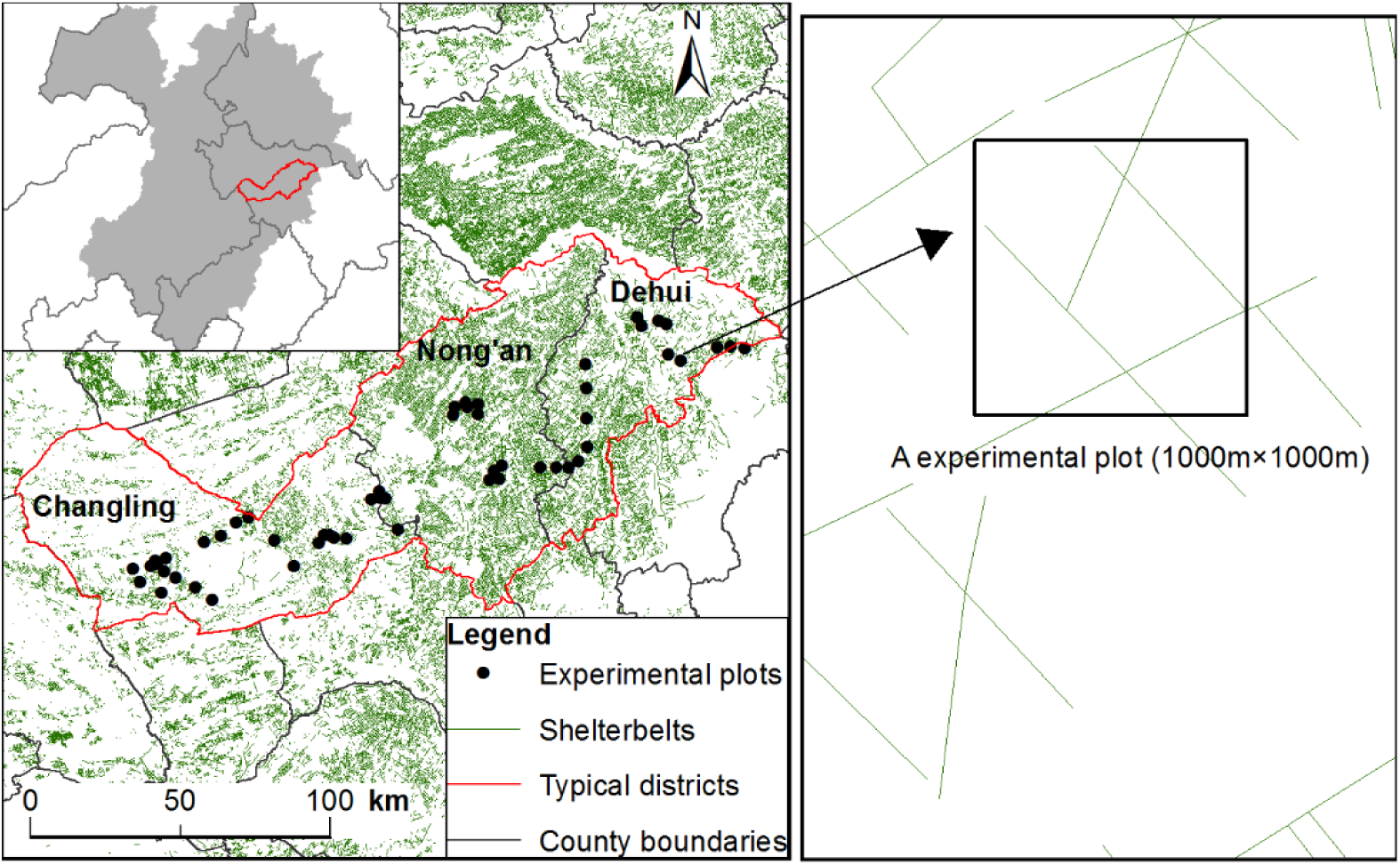
Map of maize field survey sites in Northeast China

### Monitoring of afforestation areas

#### Monitoring of forests and shrublands

Visual interpretation was used to derive the area changes of the forests and shrublands during 1978 and 2008. Although such visual interpretation is labor intensive and time consuming, the mapping accuracy of this method is higher than that of image classification using only the algorithms provided by image-processing software for the low spatial and spectral resolutions of Landsat images (6, 7). The interpretation was conducted by following the mapping principles: first, the minimum map path size was set at 6 pixels × 6 pixels, which was equivalent to 180 m ×180 m on the ground. Second, the positioning error was one pixel on the screen, which was equivalent to 30 m and 80 m on the ground in the TM/ETM+ images and the MSS images, respectively. Third, the accuracy of recognition of forests and shrublands was required to be higher than 95% according to the comparison between the result of interpretation from images (or vegetation map and topographic map) and the field survey and the statistical data from the local government. The image classification was processed using ArcMap 10.2 space analysis software (4).

The spatial resolution of TM and ETM+ images was 30 m, the area of the smallest forest or shrubland based on Landsat TM/ETM+ was 3.24 ha. While, the spatial resolution of SPOT5 was 2.5 m, the area of the smallest patch based on SPOT5 was 400 m^2^. In order to obtain more accurate forest and shrubland areas from TM images over the TNR, we therefore established correction equations in different precipitation areas (derived from TRMM data) between TM and SPOT5 images (2). The specific steps are as follows:

1) The TNR was divided into three precipitation zones, high precipitation zones (annual precipitation >= 456 mm), middle precipitation zone (annual precipitation ranging from 303 to 456 mm) and low precipitation zone (annual precipitation < 303 mm), using the Jenks natural division classification method. Precipitation was the key factor affecting the distribution of forests and shrublands in arid and semi-arid TNR (Fig. S3).
2) The correction equations on the area of forests and shrublands between SPOT and TM were obtained in the three precipitation zones. The forest areas were synchronously monitored based on TM and SPOT5 images, respectively. The typical zones (9 scenes of SPOT5 images) were divided into a number of sample units with 10000 m×10000 m (e.g., Fig. S4). Then, the statistical analysis was conducted by overlaying three maps of the climatic zones and forests and shrublands from TM and SPOT images. In each precipitation zone, we chose 80% sample units to establish the correction equations of forest areas between TM and SPOT5 (Fig. S5), and the remaining 20% sample units were used to verify the accuracy of the correction equations (Fig. S6).

**Fig. S3.**
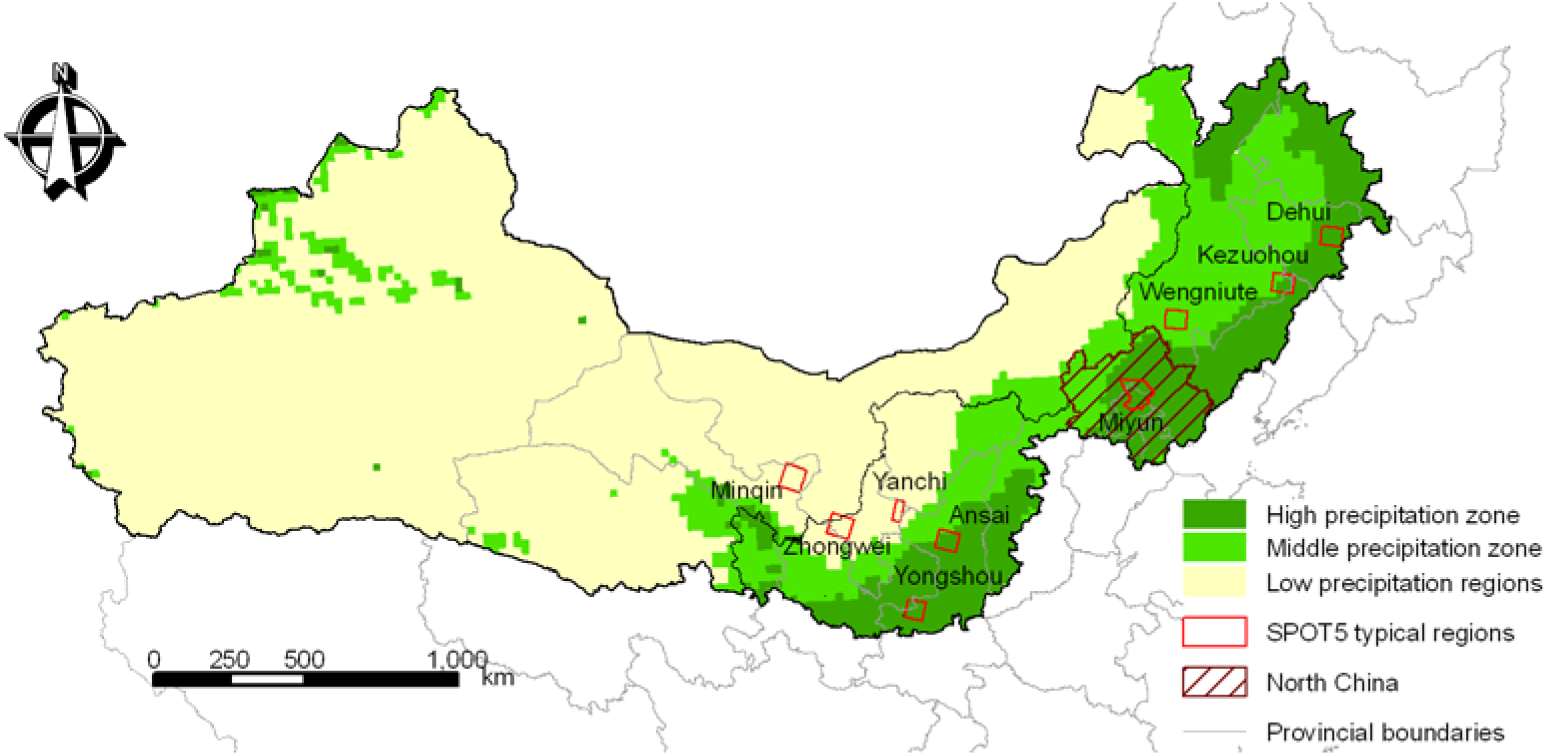
Climatic zones based on TRMM precipitation data.

**Fig. S4.**
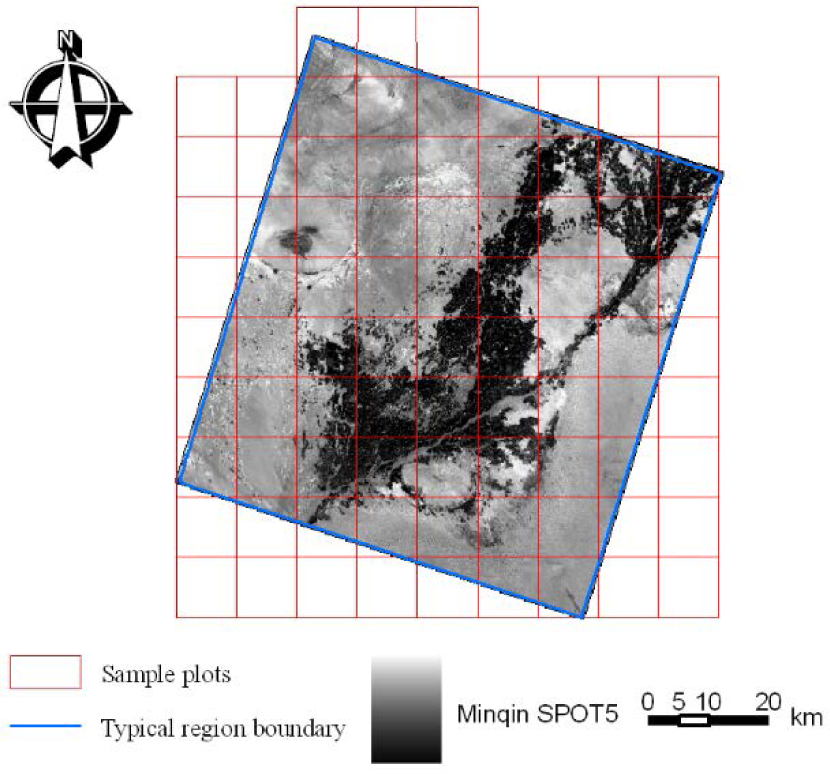
The division of plots in Minqin SPOT5 typical region.

- High precipitation zone: the high precipitation zone was located in the east and south of the TNR (Fig. S3), covering 10.92% of the total TNR, and 37.31% of forests and shrublands distributed in this zone (based on the area of forests and shrublands in 2008 estimated from TM) because of the relatively high precipitation. A total of 167 sample units were in the high precipitation zone (13845 km^2^), 134 sample units data were used to establish the correction equations (Fig. S5-a), and the remaining 33 sample units data were used to verify the accuracy (Fig. S6-a). The verification resulted in 85.4% accuracy, which was calculated as (1-RRMSE).
- High precipitation zone in the north of North China: this zone (Fig. S3) covered 2.5% of TNR, and 13.29% of forests and shrublands distributed in the region. A total of 93 sample units were in this zone (5672 km^2^), 74 sample units data were used to establish the correction equations (Fig. S5-b) and the remaining 19 sample units were used to verify the accuracy. The verification resulted in 91.1% accuracy (Fig. S6-b).
- Intermediate precipitation zone: the zone was in the middle part of TNR (Fig. S3), covering 17.99% of TNR, and 31.73% of forests and shrublands distributed in the region. A total of 90 sample units were in this zone (6235 km^2^), 72 sample units data were used to establish the correction equations (Fig. S5-c) and the remaining 18 sample units were used to verify the accuracy of the equations (Fig. S6-c). The verification resulted in 95.2% accuracy
- Low precipitation zone: this zone had the largest area, was in the western part of TNR (Fig. S3), covering 68.56% of TNR, and 17.67% of forests and shrublands distributed in the region. A total of 119 sample units were in this zone (11856 km^2^), 95 sample units data were used to establish the correction equations (Fig. S5-d) and the remaining 24 sample units were used to verify the accuracy of the equations (Fig. S6-d). The verification resulted in 72.4% accuracy

**Fig. S5.**
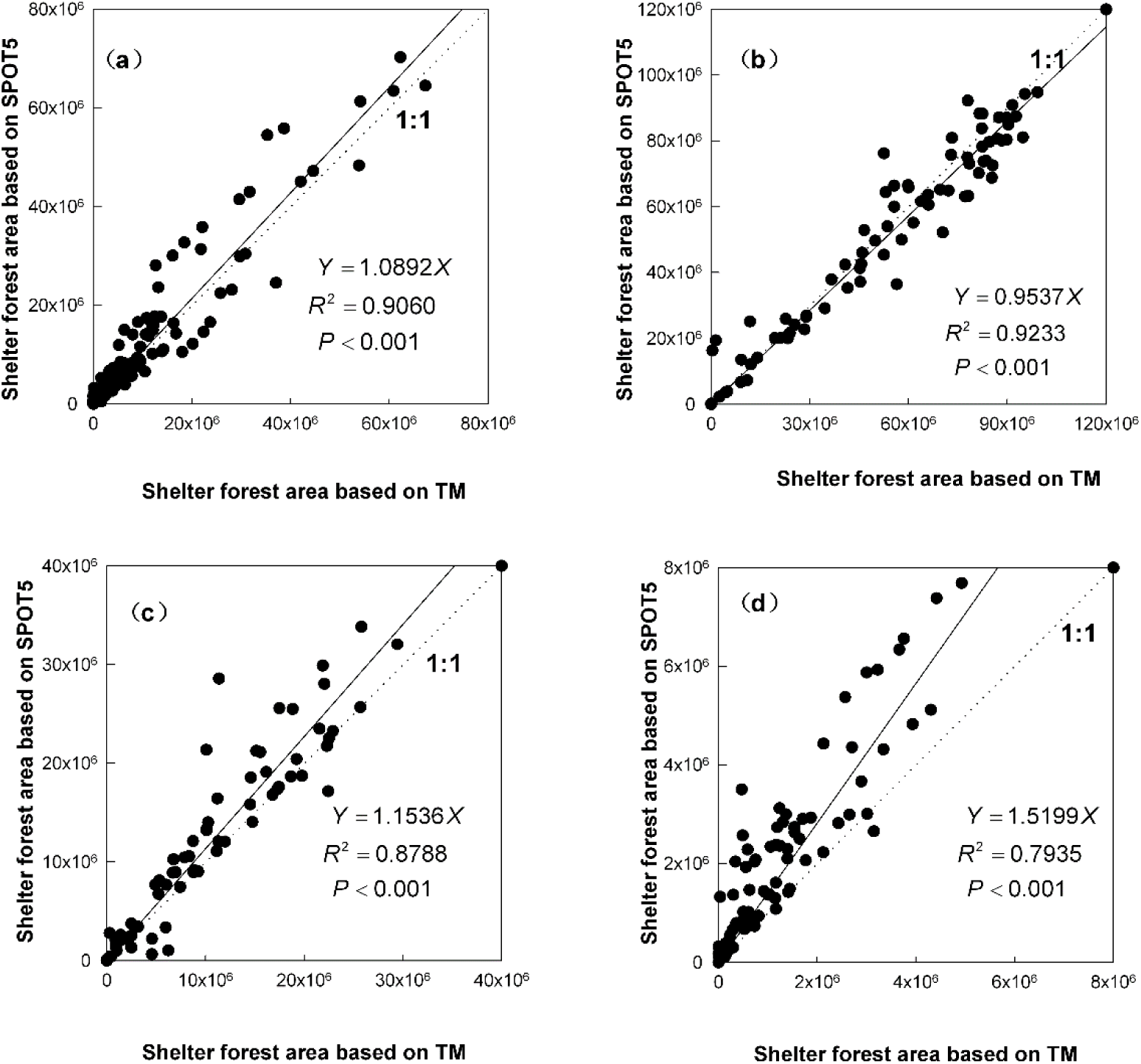
The regressive equations for the correction of forests and shrublands area in different precipitation regions derived from SPOT5 and Landsat TM: (a) high precipitation zone (n=167), (b) high precipitation zone in the north of North China (n=93), (c) Intermediate precipitation zone (n=90), and (d) low precipitation zone (n=119).

**Fig. S6.**
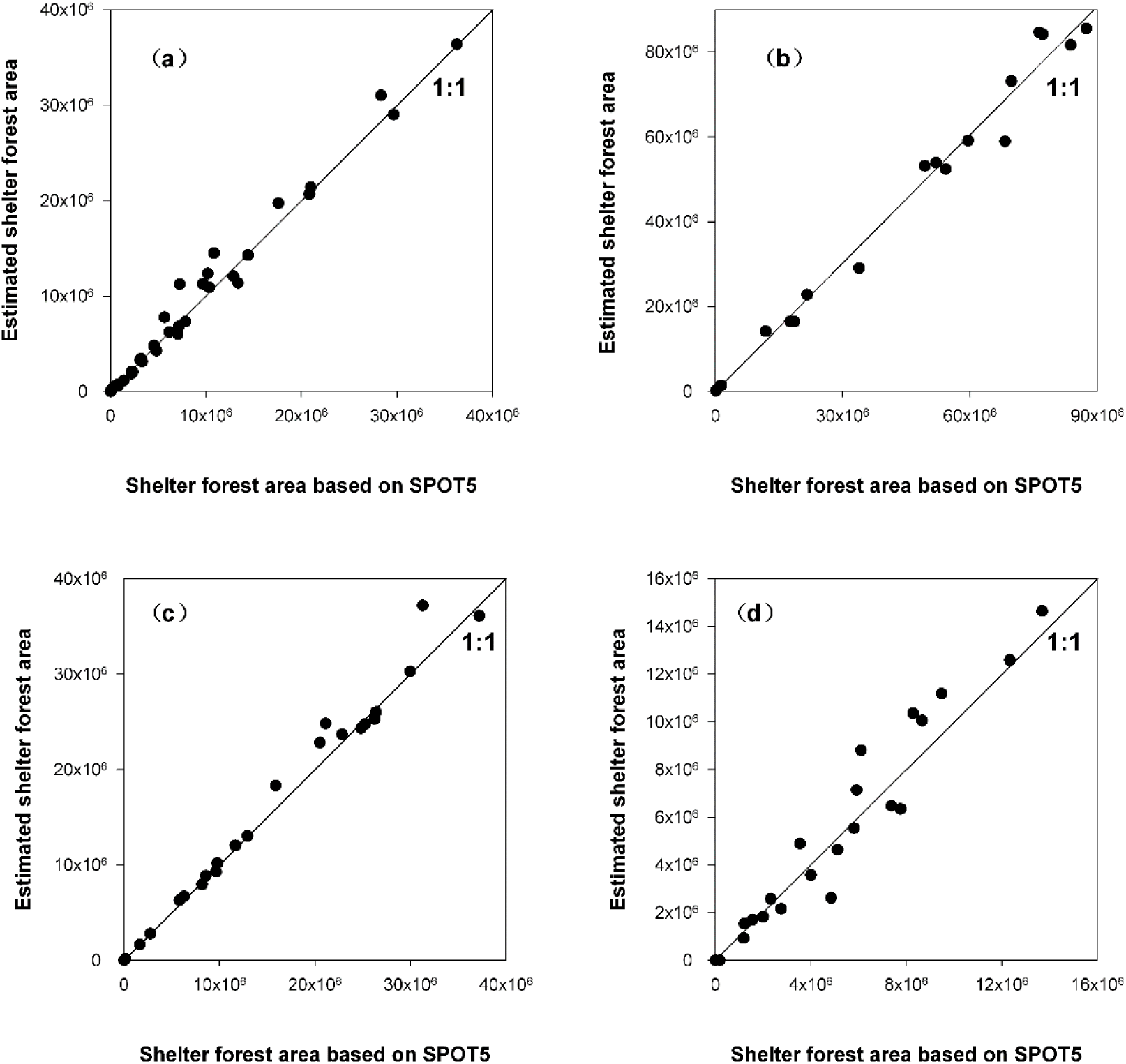
Verification of forest and shrubland area based on the relationship between SPOT5 and Landsat TM images in different precipitation regions: (a) high precipitation zone, (b) high precipitation zone in the north of North China, (c) intermediate precipitation zone, and (d) low precipitation zone.

3) The forest and shrubland areas from 1978 to 2008 under the resolution of SPOT5 were obtained. Forest and shrubland areas in 1990, 2000 and 2008 were first derived from TM Landsat images, and then corrected by the four corrected equations (Fig. S5). However the forest and shrubland areas in 1978 were derived from the Landsat MSS (the spatial resolution of 80 m), which were compared with the statistical data from the State Forestry Administration (SFA) of China. In 1978, the total area of forests and shrublands derived from the Landsat MSS was 2209.95×10^4^ ha based on remote sensing (i.e., the result in our study), while the area was 2014.49×10^4^ ha based on the statistical data from SFA. The two estimates agreed reasonably well.

#### Monitoring of shelterbelts

The shelterbelt areas were obtained by the combination of TM and SPOT5/ CBERS-02B images, and the specific steps were as follows.

##### 1) Estimation of the shelterbelt length in the farmlands of Northeast and North China

- The spatial distributions of shelterbelts in 1990, 2000 and 2008 were obtained using the visual interpretation based on Landsat imagines over Northeast and North China. In addition, the spatial distribution of shelterbelts in 2008 was obtained based on SPOT5 over the typical zone (i.e., one scene of SPOT5).
- The typical zone (i.e., one scene of SPOT5) was divided into 100 sample units, and each sample unit (7100 m ×7100 m) has the shelterbelt length data based on TM and SPOT5, respectively (Fig. S7). Among the 100 sample units, 78 were useful units, nine had no value (i.e., no shelterbelt), and 13 were abnormal.

**Fig. S7.**
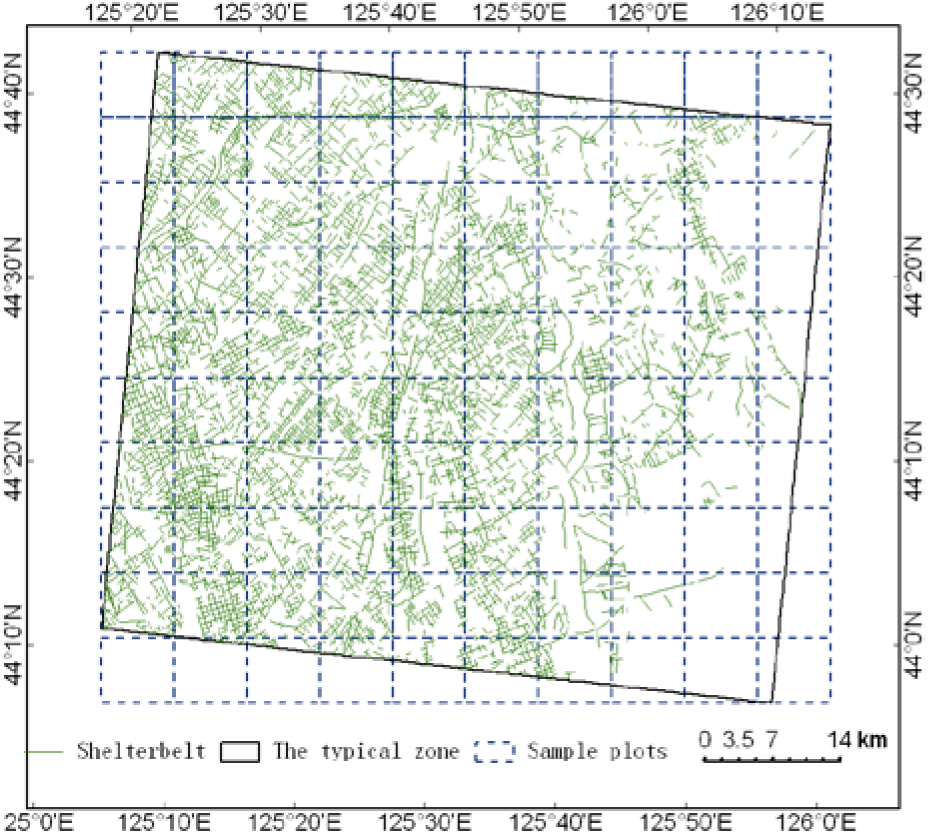
The sample units of the typical zone in Northeast and North China

- The correction relationship for the shelterbelt length between TM and SPOT5 was obtained based on the 80% of the useful sample units (i.e., 78 x 80% = 62 sample units) (Fig. S8-a), and the remaining 16 sample units were used to verify the accuracy of the correction equation (the accuracy (1-RRMSE) was about 79%) (Fig. S8-b).

**Fig. S8.**
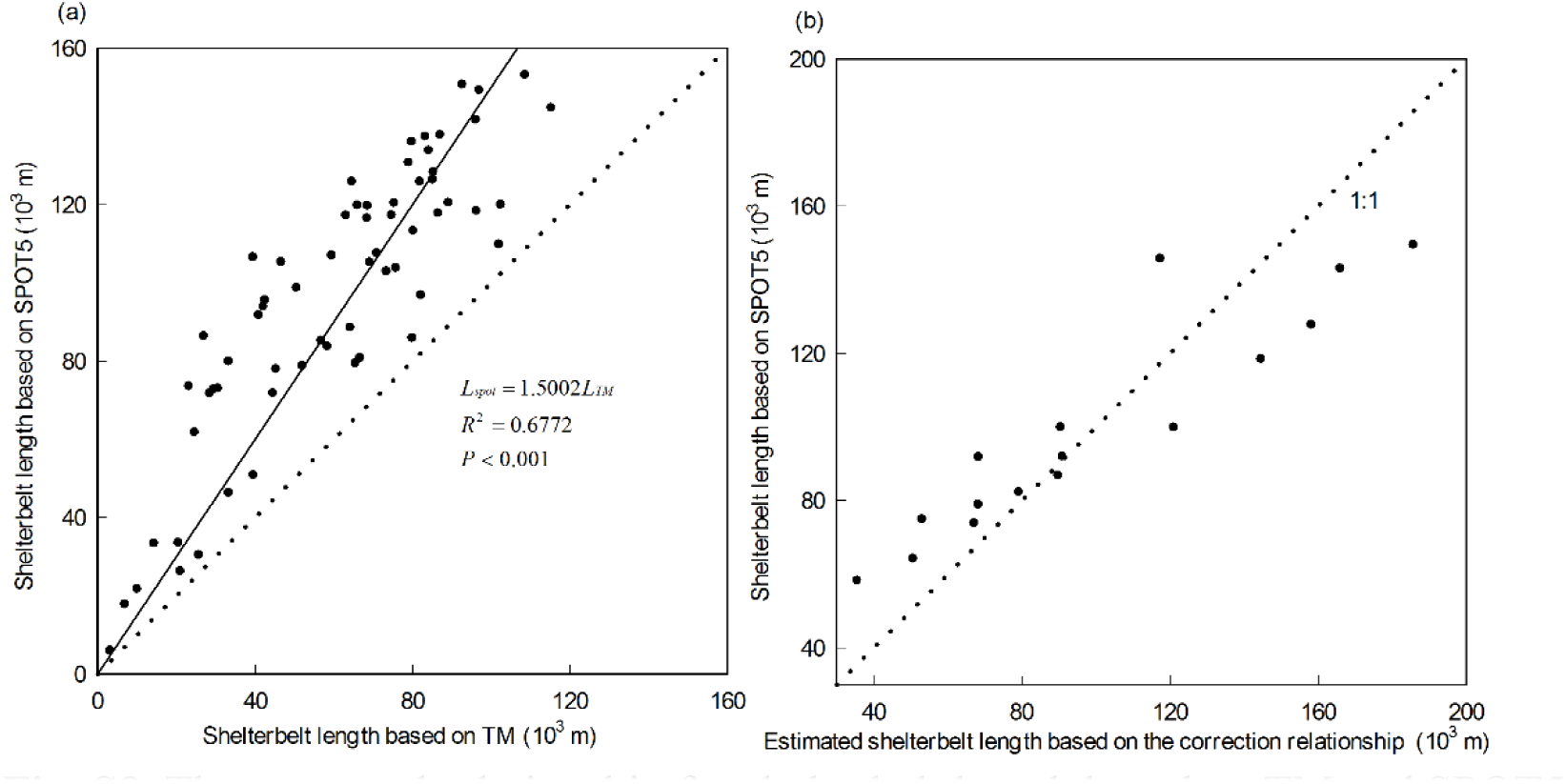
The corrected relationship for shelterbelt length based on TM and SPOT5 (a), and the corrected shelterbelt length vs. the shelterbelt length based on SPOT5 (b)

- The shelterbelt length in Northeast and North China in 1990, 2000, and 2008 was obtained by the combination of shelterbelt length data based on Landsat TM imagines (Fig. S9) and the corrected relationship between TM and SPOT5 (Fig. S8-a).

**Fig. S9.**
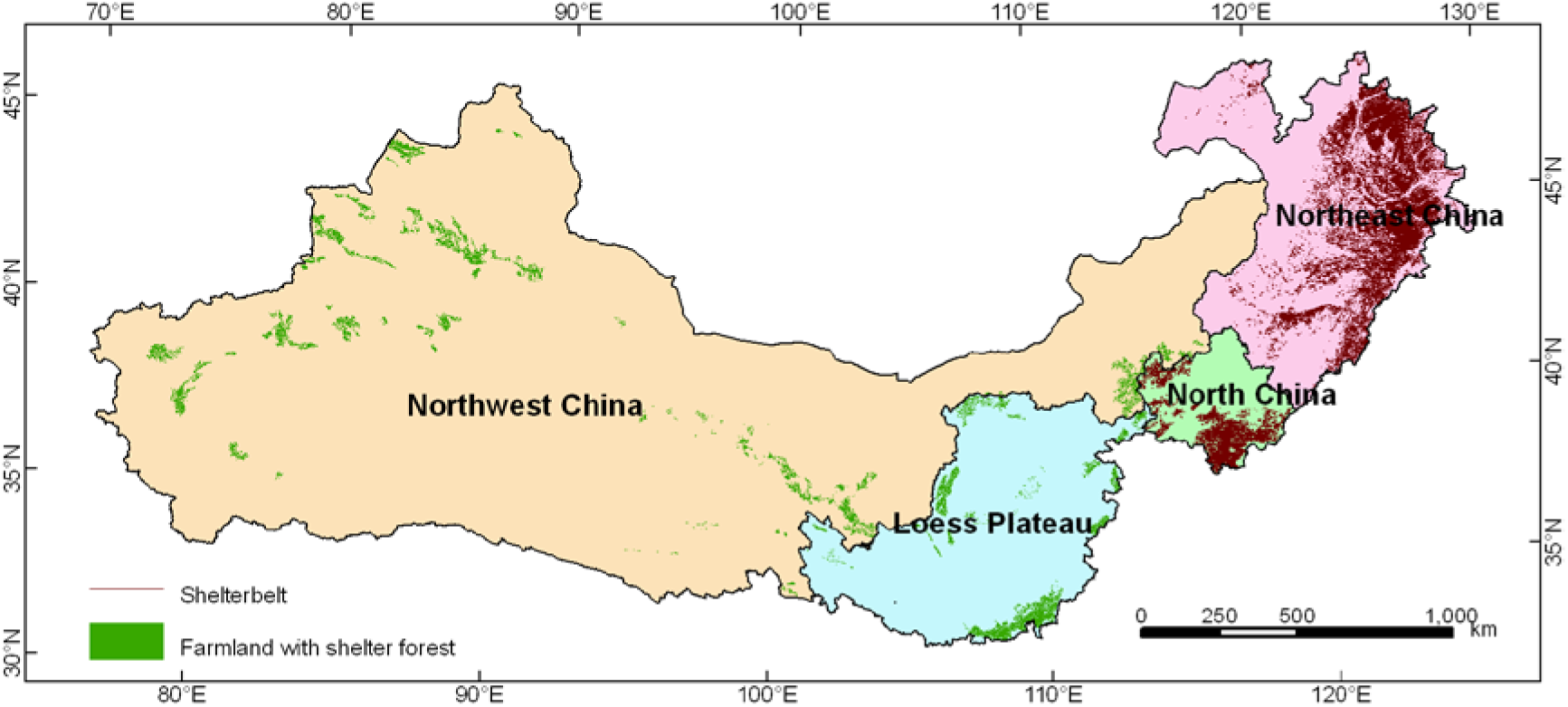
The spatial distribution of shelterbelts based on TM in Northeast and North China, and farmlands with shelterbelts based on TM in Loess Plateau and Northwest China.

##### 2) Estimates of the shelterbelt length in Loess Plateau and Northwest China

It was difficult to extract directly the shelterbelt length based on TM because the farmlands with shelterbelts were sparsely distributed in Loess Plateau and Northwest China. However, we could directly extract the farmlands with shelterbelts based on TM because farmlands were different from the other land use categories based on the distinctive color and texture in TM images data. To estimate the shelterbelt length in Loess Plateau and Northwest China, the specific processes were as follows:

- Six typical zones was chose in Loess Plateau and Northwest China, and the farmlands with shelterbelts (as polygons) were extracted based on TM, and shelterbelt length (as lines) were extracted based on CBERS-02B HR in the six typical zones (Fig. S10).
- The six typical zones were divided into a number of sample units with a 10000 m ×10000 m, and each sample unit had the spatial distribution of farmland with shelterbelt based on TM and shelterbelt based on CBERS-02B HR (Fig. S10).

**Fig. S10.**
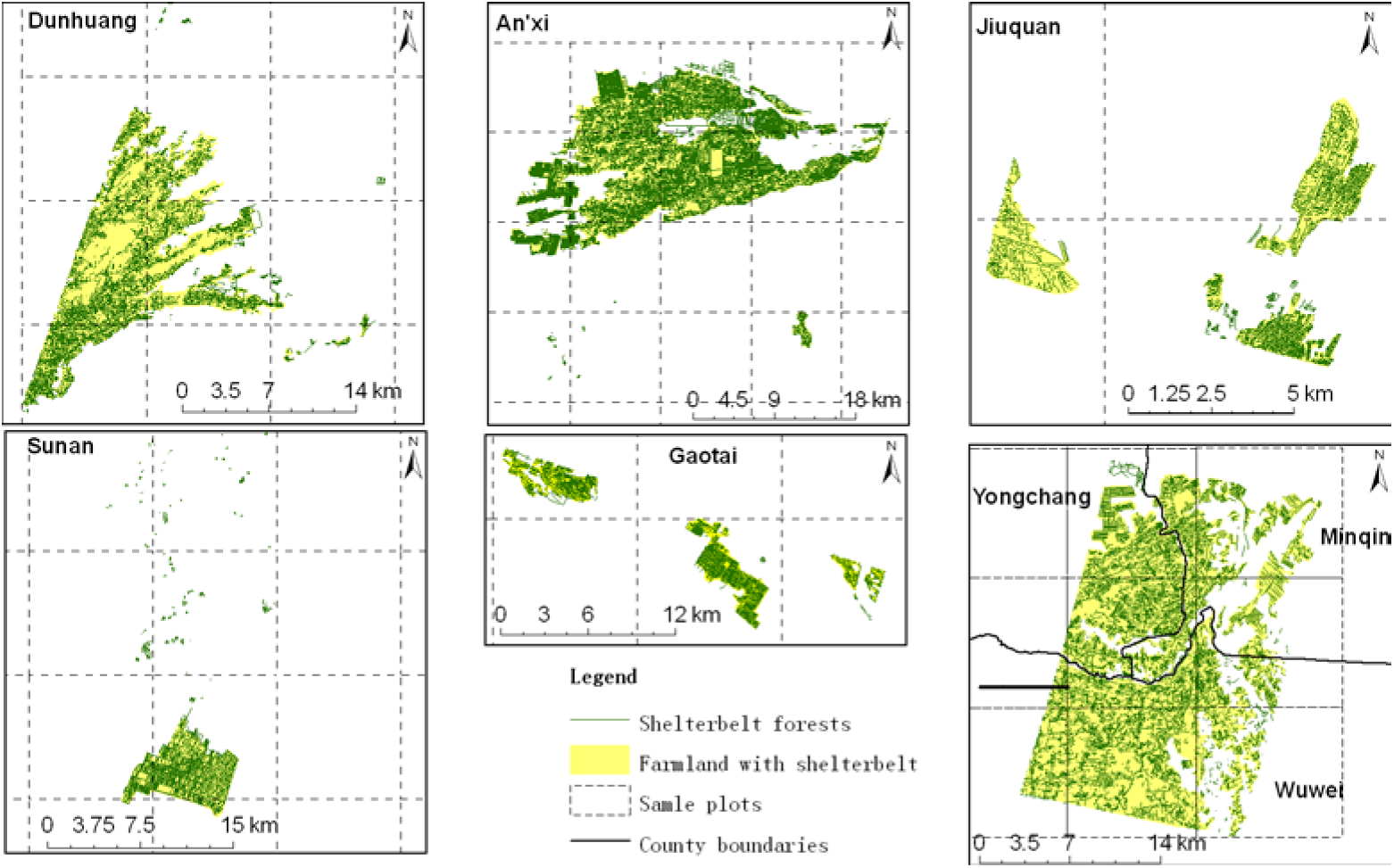
Distribution of sample plots with a 10000 m ×10000 m for farmlands with shelterbelts (based on Landsat TM) and farmland shelterbelt (based on CBERS-02B HR) in six typical zones for the Northwest and the Loess Plateau, China

- The relationship between farmland area from TM and shelterbelt length from CBERS-02B HR was established based on useful sample units. Thirty-eight sample units (10000 ×10000 m) were classified as useful units in the six typical zones (Fig. S10), among which 30 sample units (i.e., 80%) were used to build the relationship between farmland area and shelterbelt length (Fig. S11-a) and 8 sample units data were used to verify the accuracy of the correction equation (Fig. S11-b). The verification resulted in 99.80% accuracy.

**Fig. S11.**
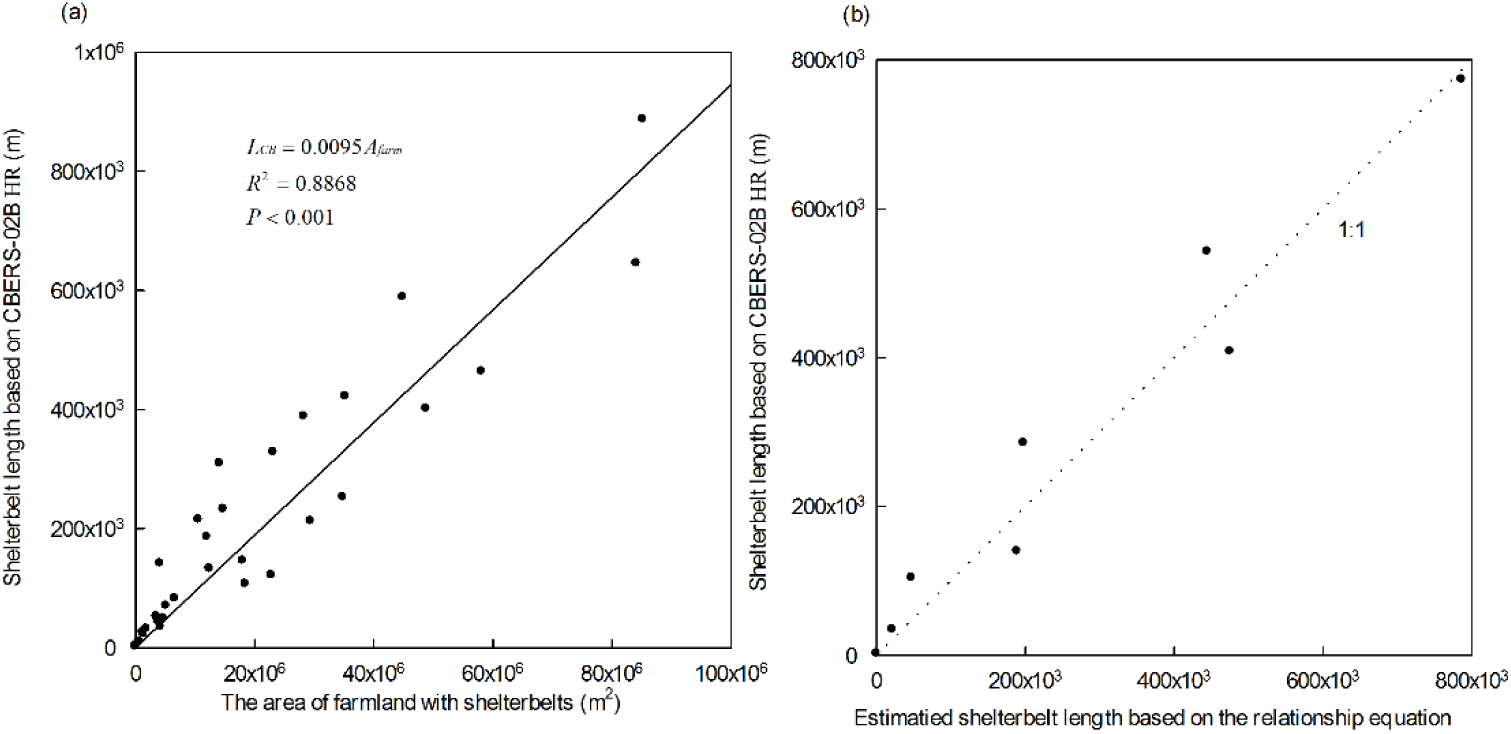
The relationship between farmland based on TM and shelterbelt length based on CBERS-02B HR (a); and the estimated shelterbelt length *VS* the shelterbelt length obtained from CBERS-02B HR (b).

- The length of shelterbelt in the Loess Plateau and Northwest China was estimated based on the above relationship equations (Fig. S11-a) and the farmland area from Landsat images (Fig. S9).

##### 3) Estimates of the shelterbelt areas in the Three-North regions

The shelterbelt area was calculated by multiplying the length of shelterbelts with their average width. The shelterbelt width was obtained on the basis of field survey. In Northeast and North China, we randomly surveyed 105 shelterbelts, including shelterbelts in farmland (accounting for 60%, the shelterbelt width was about 14 m), and shelterbelts along road and river and in other land use types (accounting for 40%, the width was about 24 m). Therefore, the weighted average width of shelterbelts was 18 m. In Loess Plateau and Northwest China, the average width of shelterbelts was 10 m according to the results of previous studies (8, 9). However, due to the limitation of spatial resolution of the Landsat MSS, the shelterbelt area in 1978 was obtained from statistical data of the State Forestry Administration of China.

#### Uncertainty

For the estimation of forested area (i.e., areas of forest and shrubland), the main uncertainties were from two sources. First, the monitoring of forested area was estimated using visual interpretation based on remotely sensed data. Although the mapping accuracy of visual interpretation method was higher than that of image classification using only the algorithms provided by image-processing software for the low spatial and spectral resolutions of Landsat images, the accuracy was about 95% based on field survey (20038 GPS points). Second, to acquire more accurate estimates of forested areas, the correction models in different precipitation areas between TM and SPOT5 images were applied. The accuracies of the correction of forested area in high precipitation zone outside the north of North China, high precipitation zone in the north of North China, intermediate precipitation zone, and low precipitation zone were 85.4%, 91.1%, 95.2% and 72.4%, respectively. According to these two sources of uncertainties, the total accuracy values of estimation of afforestation areas for forests and shrublands ranged from 68.8% (95%×72.4%) to 90.44% (95%×95.2%).

For the areas of shelterbelts, the accuracy of shelterbelts based on visual interpretation method was more than 95% by comparing with the field survey. The accuracy values of the correction procedure in the Northeast-North China and Loess Plateau-Northwest China were 79.27% and 99.80%, respectively. Therefore, the total accuracies of estimation of afforestation areas ranged from 75.3% (95%×79.27%) to 94.8% (95%×99.8%).

### Monitoring of the quality of forests and shrublands

#### Methods

The qualities of forests and shrublands were assessed using the degree of crown cover (i.e., the vegetation coverage of forests and shrublands). The vegetation coverage was estimated by the NOAA NDVI (1984-1997), SPOT NDVI (1998-1999) and MODIS NDVI (2000-2008).

For the vegetation coverage model, the principle of the dimidiate pixel model assumes that the reflectivity of each pixel can be divided into two parts (10). The reflectance values of each pixel can be defined as the weighted sum of the pure-vegetation pixel (land pixels excluding water, building and snow cover) and pure-soil pixel (non-vegetated) fractions (Equation 1):

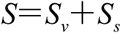

where *S* is the reflectivity of one pixel, *S_v_* is the reflectivity of the vegetative fraction of the pixel, and *S_s_* is the reflectivity of the soil fraction of the pixel (11).

Assuming the vegetation coverage of one pixel is a fraction of the vegetation coverage area (*fc*), then the ratio of the non-vegetated area to the total area is 1–*fc*. Therefore, the reflectivity of mixed pixels can be expressed as *S*=*f_c_*×*Sv*+(*1*–*f_c_*)×*S*s (12), According to the above principle, the NDVI can be approximately expressed as Equation (2),

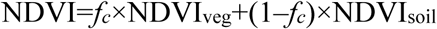

where NDVI, NDVIveg, and NDVIsoil are the NDVI values at any pixel, pure-bare pixels and pure-vegetation pixels, respectively.

Therefore, the vegetation coverage can be obtained using the following equation for a specific study area (Equation 3),

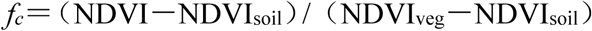

where *f_c_* is the vegetation coverage, NDVI is the value of bare-land, NDVIsoil is the minimum NDVI value in the study area, and NDVIveg is the maximum NDVI value of full vegetation cover in the study area. We identified the pure-land pixels based on the land types, the maximum and minimum NDVI values were extracted from these pixels (10).

#### Validation

We used the observed data (16 plots with 100 m×100 m) to verify the estimated vegetation coverage of forests and shrublands. The accuracy of the estimated vegetation coverage of forests and shrublands was high with R^2^ of 0.82, MAE (mean absolute error) of 2.8 mm, and RRMSE (relative root mean square error) of 6.52 %. The precision of the estimated vegetation coverage of forests and shrublands was 93.48% (1- RRMSE) (Fig. S12).

**Fig. S12.**
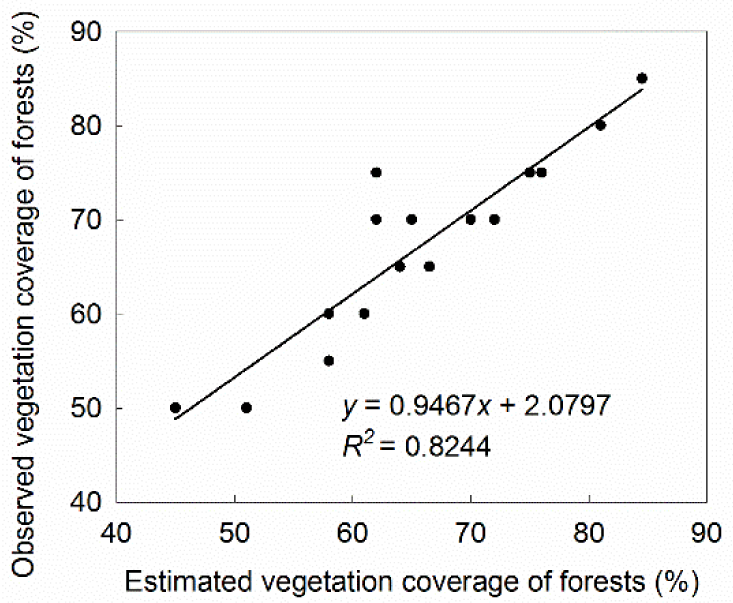
Estimated vegetation coverage based on remote-sensing VS. ground survey measurements.

### Impact on crop yields

#### 1) Climatic potential crop productivity

The climatic potential productivity of maize was calculated using the following equation (Equation 4),

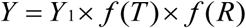

where *Y*_1_ is the photosynthesis potential productivity, *f* (*T*) is the temperature modification function, and *f* (*R*) is the water correction function.

The parameter *Y*_1_ can be calculated as Equation (5),

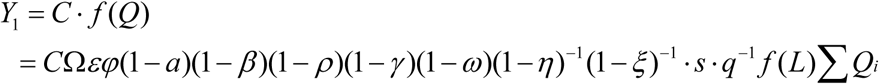

where C is the unit conversion coefficient, and *Q_i_* is the total radiation of the sun (MJ m^-2^). The spatial and temporal characteristics of *Q_i_*, which is a function of latitude that assumes a planar land surface, depend on the Julian date and latitude, and can be computed as described by Allen, Pereira (13). Definitions of other parameters are given in **Table S1**.

**Table S1.** Meaning and value of parameters used to calculate maize photosynthesis potential productivity

| Parameter | Physical meaning | Value |
| --- | --- | --- |
| $\varepsilon$ | The ratio of photosynthetic radiation to total radiation | 0.49 |
| $\varphi$ | Quantum efficiency of photosynthesis | 0.224 |
| $\alpha$ | Plant population reflectivity | 0.68 |
| $\beta$ | Plant rankness population transmittance | 0.06 |
| $\rho$ | Ratio of radiation intercepted by non-photosynthetic organs | 0.10 |
| $\gamma$ | Ratio of light over the saturation point | 0.01 |
| $\omega$ | Ratio of respiration consumption to photosynthetic product | 0.30 |
| $\eta$ | Water content of mature corn | 0.15 |
| $\zeta$ | Ratio of plant abio-ash contents | 0.08 |
| $s$ | Crop economic coefficient | 0.40 |
| $q$ | Heat content of dry matter (MJ kg<sup>-1</sup>) | 17.20 |
| $\Omega$ | Ratio of CO<sub>2</sub> photosyntheticly fixated by crop | 1.00 |
| $F(L)$ | Modification value for the dynamic change of leaf area | 0.58 |

The function *f* (*T*) can be calculated as:

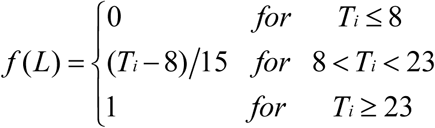

where T is the mean monthly temperature during the growing season (i.e., from May to October in this study).

The function *f* (*R*) is the water correction function for crop growth, development and yield, and can be calculated as Equation (7),

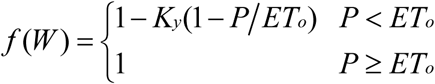

where *Ky* is the crop coefficient (i.e., 1.25 for maize in this study); *ETo* is the crop reference evaporation, which describes the water required by a given crop; and *P* is the total precipitation.

The average mean monthly air temperature, crop reference evaporation and precipitation during the growth period (i.e., from May to October) from 2007 to 2009 were estimated using MODIS and TRMM 3B43 data (1, 14, 15).

#### 2) The level of protection

To quantify the level of protection on farmland provided by the shelterbelts at the regional scale, approximately 121,011 km^2^ farmland was identified from Landsat TM images taken in 2008 using the visual interpretation method (4). The farmland was divided into grids with a 1-km resolution that were consistent with MODIS data sources (e.g., climatic potential crop productivity) in this study. The level of protection of each grid of farmland was then computed. Previous studies showed that the level of protection depended on the effectively protected distance and the growth condition (i.e., quality) provided by the shelterbelts (16–18). Therefore, the level of protection of each grid of farmland was calculated with the following formula (Equation 8),

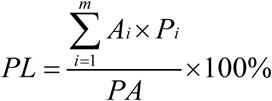

where *PL* is the level of protection of each grid of farmland; *A_i_* is the maximum area of effective protection provided by the shelterbelts in grid net *i*, which was derived from the distance of effective protection for each shelterbelt; *P_i_* is the quality coefficient of each shelterbelt in grid net *i*; and *PA* is the pixel area (i.e., 1000000, 1 km × 1 km, the spatial resolution of the objective).

##### (1) Maximum area of effective protection

The maximum area of effective protection is defined as the maximum sheltered zone (i.e., the region of reduced wind speed) that lies within the effectively protected distance of the shelterbelts (17, 19). The crop yield increase due to shelterbelts was found to vary with climatic conditions and locations. The reduction in the relative wind speed behind relatively narrow, homogenous tree windbreaks was measured, and was used to investigate the effects of angles between a given shelterbelt and the wind direction with regard to the crop yield increase due to the shelterbelt (17, 18).

The maximum effectively protected distance of each shelterbelt ranged from 15 times of shelterbelt height (i.e., 15 H, where the shelterbelt is parallel to the direction of the prevailing winds) to 25 times of shelterbelt height (i.e., 25 H, where the shelterbelt is perpendicular to the direction of the prevailing wind) according to a series of field experiments over 15 years in Northeast China (17, 18). Therefore, 20 H, which is the average of 25 H and 15 H, was chosen as the maximum effectively protected distance for each shelterbelt.

The shelterbelt height is a critical factor determining the extent of the protected area of a given shelterbelt. *Populus* spp. (including *P. canadensis, P. beijingensis, P. xiaozuanrica, P. simonii, and P. pseudo-simonii*) are the primary tree species used in shelterbelts and account for more than 90% of shelterbelts in Northeast China (20). Based on previous studies, each shelterbelt height was related to the stand age (17), whereas the shelterbelt age could be obtained using remote-sensing technologies. Based on the fundamentals of phase-directional management for shelterbelts (20), the shelterbelt ages were categorized as < 10, 10-20, and > 20 years. The procedures used to estimate the shelterbelt ages based on a time series of Landsat images are described as follows.

• Extract the spatial distribution of shelterbelts in 1990, 2000 and 2008

A time series of Landsat TM and ETM+ images were used to determine the changes of shelterbelts. First, in order to guarantee to have a clear image, only the images with cloud cover < 10% were selected. For interpreting the shelterbelt data, images in early May or late October were used to distinguish shelterbelts from crops in 1990, 2000 and 2008. Second, the 2000 ETM+ images were georeferenced and orthorectified using 50 to 60 ground control points (GCPs) that were derived from 1:100,000 topographic maps. The RMSE for the geometric rectification was less than 1 pixel (i.e., < 30 m). TM images from 1990 and 2008 were then matched with the 2000 ETM+ images using the mean of the image-to-image matching method. During the image-matching process, 40 to 50 GCPs were randomly selected in the images from 1990 and 2000 to cover most of the area represented by the two sets of images. The RMSE of the geometric rectification between the two images was 1 to 2 pixels (i.e., < 60 m) in flat areas and 2 to 3 pixels (i.e., < 90 m) in mountainous areas. Finally, the corrected images were used to interpret the shelterbelt data from 1990, 2000 and 2008 using the visual interpretation method.

• Determination of shelterbelt ages in 2008

The determination of shelterbelt ages was conducted in the following list.

If the shelterbelts did not appear in the ETM+ images in 2000, but appeared in the TM images in 2008, the associated shelterbelts were assumed to have been planted between 2000 and 2008; thus, the age of these shelterbelts should be less than 10 years.

If the shelterbelts did not appeared in the TM images in 1990, but appeared in the ETM+ images in 2000 and in the TM images in 2008, then, these shelterbelts were assumed to have been planted between 1990 and 2000; thus, the age of the shelterbelts was 10 to 20 years.

If the shelterbelts appeared in the TM images in 1990, the ETM+ images in 2000 and the TM images in 2008, these shelterbelts were assumed to have been planted between 1978 and 1990, and the age of shelterbelts was more than 20 years.

According to the ages assigned above, the spatial distribution of the shelterbelt ages in 2008 was derived based on the spatial distribution of shelterbelts in 1990, 2000 and 2008 using ArcGIS software.

##### (2) Quality coefficient of shelterbelts

The quality of shelterbelts (e.g., optical porosity) has significant effects on the relative wind speed reduction across a given landscape (16, 18). The normalized difference vegetation index (NDVI) is an index of plant “greenness” or photosynthetic activity, and is one of the most commonly used vegetation indices. Researchers have used the NDVI to assess large-scale patterns in vegetation quality, net primary productivity, and biomass with mixed results (21–23). In this study, NDVI was used as a proxy for the quality of shelterbelts, and to determine the quality (i.e., growth conditions) coefficient of the shelterbelts based on the Landsat TM images.

The quality coefficients of different age groups based on NDVI values were calculated as follows. First, NDVI values across Northeast China were calculated from Landsat TM images taken in 2008. Second, the mean NDVI value of each shelterbelt was extracted using a combination of shelterbelts and NDVI values. Third, in different shelterbelt age groups, the mean (*u*) value and standard deviation (*σ*) of the NDVI were calculated. The different shelterbelt age groups were also divided into good (NDVI ≥ *u+σ/2*), normal (*u-σ/2* ≤ NDVI < *u+σ/2*), and poor (NDVI < *u-σ/2*) quality. Fourth, the mean NDVI values in different shelterbelt age groups were also calculated. Fifth, we hypothesized that shelterbelts within different age groups could produce the maximum effectively protected distance (i.e., 20 H) if they were of good quality; therefore, quality coefficients were defined as 100% for good quality (i.e., NDVI ≥ *u+σ/2*) in all age groups. For normal and poor quality shelterbelts of different age groups, the quality coefficients were defined as the ratio of mean NDVI values in the normal and poor quality to that in good quality, respectively. Finally, the quality coefficients were obtained for the different shelterbelt age groups.

#### 3) Estimation of maize yields

The harvest index (HI) was applied to estimate the maize yields based on MODIS gross primary productivity (GPP) data. The parameter HI, which is the ratio of grain yield to total the aboveground biomass, is a key parameter when predicting crop yields using remote sensing data (24). The MODIS GPP is a cumulative composite of GPP values based on the radiation-use efficiency concept that provides a new opportunity for timely wheat yield estimates at the regional scale (25). The maize yields were estimated because they accounted for more than 50% of the farmland in Northeast China in this study.

Annual composite MODIS GPP data (MOD17A3) at a 1-km spatial resolution from 2007 to 2009 were downloaded from the Earth Observing System Gateway (online at https://lpdaac.usgs.gov/). GPP estimates from the MODIS data are reported in kg C m^-2^, which are easily converted into biomass estimates because carbon comprises approximately 50% of vegetative biomass (26). At physiological maturity, approximately 90% of the accumulated biomass of maize is aboveground, whereas the remainder is located in the plant’s roots (25). The harvest index method was calculated for regional-scale maize yields using the following equation (Equation 9),

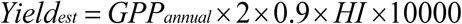

where *Yield_est_* is the estimated yield (kg ha^-1^); *GPP_annual_* is the average annual GPP from 2007 to 2009, which is derived from the MODIS GPP data (kg C m^-2^); 2 is a conversion factor from carbon (kg C m^-2^) to biomass (kg C m^-2^); 0.9 (i.e., 90%) is the annual proportion of GPP that is allocated to aboveground productivity; *HI* is the harvest index, which was set to 0.703 by comparing 17 county-level maize yields data in 2008 from the China Statistical Yearbook, and the values of *GPP_annual_*×2×0.9; and 10000 is a unit conversion factor from kg C m^-2^ to kg ha^-1^.

To validate the estimation of maize yields at a 1-km resolution at the regional scale, 58 experimental farmland plots that were 1000 m × 1000 m were collected in a typical district, including the counties of Dehui, Nong'an, and Changling in Northeast China in 2008. The coefficient of determination (R^2^) and the root mean square error (RMSE) were used to determine the accuracy of the estimated maize yields in Northeast China (*R*^*2*^=0.74, RMSE=520.02 kg ha^-1^).

#### 4) The contribution rate to increase crop yields

First, to reduce the influence of climatic factors on crop yields, the farmland was divided into high, medium, and low climatic potential productivity based on the climatic potential productivity of maize. Second, the method of the level of protection at regional scales was created and maize yields were estimated by using the remote sensing data. Third, maps that describe the level of protection, maize yields and climatic potential productivity zones were created, where all values were in pixels, and all pixel values of the level of protection and the corresponding maize yields were determined under the different climatic potential productivity zones (i.e., high, medium, and low potential productivity). Fourth, farmland in the different protection levels in the different climatic potential productivity zones included nearly all soil types and microclimate conditions; thus, the averaging method at the pixel scale can be used to determine the crop yield in the different protection levels in the mean soil type and microclimate conditions. The level of protection was the only factor considered in this study; we averaged the corresponding maize yields (i.e., the average values of maize yields under different soil types and microclimate conditions) in all grids in the different climatic potential productivity zones, which effectively reduced the influence of these factors. Thus, all pixels of protection level 0 (i.e., unsheltered fields) and the different protection levels (i.e., 10%, 20%, 30%, …, 90%, and 100%) and the corresponding maize yields in the different climatic potential productivity zones were determined. We then computed the average crop yields under different protection levels (i.e., 0%, 10%, 20%, 30%, …, 90% and 100%). The relationship between the average protection level and the corresponding maize yields in the different climatic potential productivity zones was then determined. Finally, we analyzed the effects of the shelterbelts on the maize yields and computed the contribution rates of shelterbelts with regard to increasing maize yields in the different climatic potential productivity zones at the regional scale. The contribution rate was calculated as Equation (10),

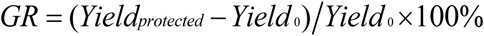

where *GR* is the contribution rate of shelterbelts to increase maize yield, *Yield_protected_* is the maize yield at a given protection level (kg ha^-1^), and *Yield*0 is the average maize yields (kg ha^-1^) at protection 0, which indicated an unsheltered field (5). The greatest contribution rate would occur when all farmland were under optimal level of protection (>80%) from shelterbelts.

### Impact on soil erosion by water

#### 1) Estimation of soil erosion by water

The average soil loss (A) due to water erosion per unit area per year was quantified using Revised Universal Soil Loss Equation (RUSLE) (27), which is an empirically based model founded on the Universal Soil Loss Equation (USLE) (28) and has been described in detail by Renard et al. (29). The RUSLE is expressed as:

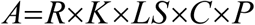

where *A* is the estimate of average annual soil loss (Mg km^-2^ year^-1^) caused by water erosion; *R* is the rainfall erosivity factor to reflect rainfall pattern (MJ mm ha^-1^ h^-1^ year^-1^); *K* is the soil erodibility factor to indicate soil type (Mg ha h ha^-1^ MJ^-1^ mm^-1^), which is a measure of the susceptibility of soil to be eroded under standard conditions; *LS* is the topographic factor, derived from a combination of the slope steepness and slope length measurements (non dimensional); *C* is the vegetation cover and management factor (non dimensional); *P* is the support practice factor (non dimensional). Each factor in *RUSLE* model was calculated using ArcGIS, and multiplying factor map layers in the ArcGIS gave the spatial distribution of the soil loss of the TNAP.

##### • Calculation of *R*

Rainfall erosivity factor (*R*): the values used for the rainfall erosivity factor (*R*) in RUSLE quantify the effect of raindrop impact, and reflect the amount and rate of runoff likely to be associated with the rain (27).

The *R* factor for any given period is obtained by summing for each rainstorm— the product of total storm energy (*E*) and the maximum 30-min intensity (*I*30). Since pluviograph and detailed rainstorm data are rarely available at standard meteorological stations, mean annual (30, 31) and monthly rainfall amount (32) have often been used to estimate the *R* factor for the RUSLE. In an effort to estimate the *R* factor using monthly and annual rainfall data, Wischmeier and Smith (28) proposed an empirical formula for estimating the *R* value, which is defined as equation (12),

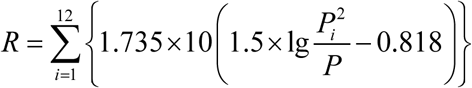

where *P*_i_ is the mean rainfall amount in mm for month *i*; *P* is the rainfall total for the same year amount in mm.

There are 220 rain gauges in the Three-North regions used in this study. These data were distributed in monthly mean rainfall. The *R* factor was calculated using equation (11) for each meteorological station. The rainfall data in 1980 were used to calculate the *R* factor from 1978. The rainfall data from 1981 to 1990, from 1991 to 2000, from 2001to 2008, were used to calculate the *R* factors of 1990, 2000, and 2008, respectively. Inverse Distance Weighting (IDW) interpolation was applied to create rainfall erosivity maps for 1978, 1990, 2000 and 2008 for the study region.

##### • Calculation of *K*

Soil erodibility factor (*K*): The K factor is an empirical measure of soil erodibility as affected by intrinsic soil properties (33). The main soil properties affecting *K* include soil texture, organic matter, structure, and permeability of the soil profile. Soil erodibility (*K* factor) was estimated with the help of the soil map provided by the second national soil census of China during 1979 and 1994 at a scale of 1:1,000,000, which is available online on http://www.geodata.cn/Portal/. The spatial distribution maps of soil types were obtained with the help of ArcGIS. Afterwards, the K factor was calculated by Equation (13), which were developed from data of measured K values (27).

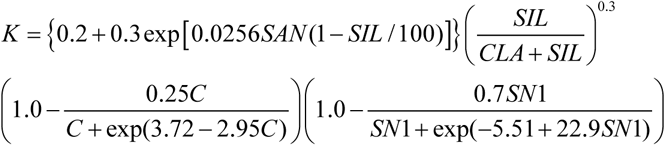

where *SAN, SIL, CLA*, and *C* are the content of sand, silt, clay and soil organic carbon, respectively; *SN*1=1-*SAN*/100. Table S2 gives an overview of the texture parameters and estimated *K* values for each texture class.

Four periods of *K* factor were obtained by the same distribution map of *K* factor.

The mean *K* value was 0.0594 Mg ha h ha^-1^ MJ^-1^ mm^-1^, the maximum value was 0.0940 Mg ha h ha^-1^ MJ^-1^ mm^-1^, which mainly distributed in the Loess Plateau and the North China.

**Table S2.** Representative soil texture parameters for each texture class

| Dominant soil textural class | % clay | % silt | % sand | K values |
| --- | --- | --- | --- | --- |
| Coarse (clay < 18% and sand > 65%) | 9 | 8 | 83 | 0.0115 |
| Medium (18% < clay < 35% and sand > 15%, or clay < 18% and 15% < sand < 65%) | 27 | 15 | 58 | 0.0311 |
| Medium fine (clay < 35 % and sand < 15 %) | 18 | 74 | 8 | 0.0438 |
| Fine (35 % < clay < 60 %) | 48 | 48 | 4 | 0.0339 |
| Very fine (clay > 60 %) | 80 | 20 | 0 | 0.0170 |

##### • Calculation of *LS*

The topographic factor (*LS*) reflects the effect of topography on erosion by water. It has been demonstrated that the increasing of slope length and slope steepness can produce higher overland flow velocities and correspondingly higher erosion (34). Moreover, gross soil loss is considerably more sensitive to changes in slope steepness than to changes in slope length (35). Slope length has been broadly defined as the distance from the point of origin of overland flow to the point where either the slope gradient decreases enough where deposition begins or the flow is concentrated in a defined channel (28). Two different parameters are used to calculate the LS-factor, flow length and flow accumulation in this study. With the help of ArcGIS, the original DEM with 90 m resolution was firstly converted to slope map in degree and flow direction map. Afterwards, the flow direction map was used to create maps of flow length and flow accumulation. The *LS* factor was estimated according to Moore and Burch (36) (Equation 14).

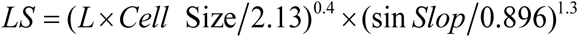

where L is flow length; Cell Size is the size of pixel; Slop is the slope of terrain.

##### • Calculation of *C*

The Cover and Management factor (*C*): the vegetation cover management factor *C* represents the effects of plants, crop sequence and other cover surface on soil erosion by water. The value of *C* is defined as the ratio of soil loss from a certain kinds of land surface cover conditions (28).

NDVI can be used as an indicator of the land vegetation vigor and health (37). In addition, the satellite remote sensing data could act as an extremely important role to estimate the C factor due to the variety of the land cover patterns (38, 39). Therefore, the relationship between NDVI and *C* values was used to obtain the *C* factor in this study. NDVI derived from the original satellites of NOAA/AVHRR from 1980 to 1999, and from MODIS for 2000 to 2008, were used to calculate the annual *C* factor by applying the relationship used in *(g40)* and (41):

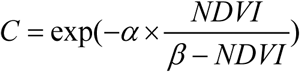

where *C* is the calculated cover management factor; NDVI is the vegetation index, and *α* and *β* are two scaling factors. The values for the two scaling factors *α* and *β* were given as 2 and 1, respectively (42).

The *C* values of 1980 were represented as those of 1978, and the mean annual *C* values from 1981 to 1990, from 1991 to 2000, and from 2001 to 2008 represented those of 1990, 2000 and 2008, respectively.

##### Calculation of *P*

The conservation practice factor (*P*): The conservation practice factor (*P*) is also called as support factor. It represents the soil-loss ratio after performing a specific support practice to the corresponding soil erosion by water, which can be treated as the factor to represent the effect of soil and water conservation practices (27, 43). *P* ranges from 0 to 1. The lower the value is, the more effective the conservation practices are. The *P* values for common support practices were obtained from experimental data under runoff–erosion plots with different support practices using both natural and simulated rainfalls (9). However, at a large watershed scale, it is very difficult or impossible to measure the *P* factor of every plot. Therefore, only a rough *P* factor value was calculated using the Wener method (33, 44) (Equation 16).

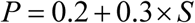

where *S* is the slope grade (%), which was derived from the DEM.

#### 2) Categorization of soil erosion levels

The soil loss by water was classified into soil erosion maps with five different soil erosion levels according to the Standards for Classification and Gradation of Soil Erosion (The Ministry of Water Resources of the People’s Republic of China, 2009). The threshold for each of the soil erosion levels is presented in **Table S3**.

**Table S3.** Classification of soil erosion intensity in the regions of the Three North Forestation Program.

| Levels | Average soil erosion modulus<br>(Mg km<sup>-2</sup> year<sup>-1</sup>) |  | Average erosion depth (mm year<sup>-1</sup>) |  |
| --- | --- | --- | --- | --- |
|  | Loess region | Other Regions | Loess region | Other Regions |
| without | <1000 | <200 | <0.74 | <0.15 |
| slight | 1000-2500 | 200-2500 | 0.74-1.9 | 0.15-1.9 |
| moderate | 2500-5000 | 2500-5000 | 1.9-3.7 | 1.9-3.7 |
| severe | 5000-8000 | 5000-8000 | 3.7-5.9 | 3.7-5.9 |
| very severe | 8000-15000 | 8000-15000 | 5.9-11.1 | 5.9-11.1 |
| extremely<br>severe | >15000 | >15000 | >11.1 | >11.1 |

In additions, to verify the precision of the estimation of soil erosion by water based on the *RUSLE*, more than 20 ground-based observations in the Loess Plateau from 2004 to 2007 were used, including 16 counties and four watersheds (Yahe River, Qingshuigou watershed, Majiagou watershed, and Qingshui River). These datasets were obtained from the hydrological stations within the counties or watersheds. The average soil erosion modulus in 2008 was extracted for the 16 counties and the four watersheds. The accuracy of the results based on the *RUSLE* were compared with the observation by paired *t*-test (*P*＝0.075).

#### 3) Path analysis to determine the contribution rate to soil erosion by water

Path analysis was applied to determine causal relationships between the coverage of forests or shrublands, and the processes of soil erosion by water (45. The direct and indirect effects in the path analysis were derived from (i) multiple linear regression of environmental and vegetation factors on the processes of soil erosion by water, and (ii) simple correlation coefficients between possible factors influencing the erosion. The direct effects of the factors influencing the processes of soil erosion by water were termed path coefficients, and we computed standardized partial regression coefficients for each of the factors in the multiple linear regression against the processes of soil erosion by water (46). The indirect effects of the factors influencing the processes of soil erosion by water were determined from the simple correlation coefficient between the factors and the path coefficients. The correlation between the processes of soil erosion by water and any one factor was the sum of the entire path connecting two variables, as described by Equation (17).

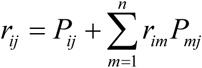

where *r*_*ij*_ is the simple correlation coefficient between the processes of soil erosion by water and a factor influencing the erosion; *P*_*ij*_ is the path coefficient between the processes of soil erosion by water and any one factor influencing the erosion; and *r*_*im*_*P*_*mj*_ is the indirect effect of any one factor influencing the processes of soil erosion by water.

The seven possible factors influencing erosion included precipitation, evapotranspiration, amount of coniferous forests, amount of broadleaved forests, amount of mixed forests, amount of shrublands, and the total vegetation cover resulted from both water conditions and the establishment of forests and shurblands. The stepwise regression procedure was applied in the present study to eliminate the independent variables (the factors influencing the erosion) that do not contribute significantly to the fit of the regression model. Only statistically significant variables (*P*<0.05) were kept in the regression. *R*^2^ (the coefficient of determination of the multiple regression equation) was used to evaluated the efficiency of the regression.

Pearson correlation analysis was performed for the five types of soil erosion by water (very slight, slight, moderate, severe and extremely severe soil erosion) among the factors influencing the erosion at 5% level of significance (Table S4). These statistical analyses were performed using SPSS 16^th^ Edition (Chicago, USA). If significant correlation was found between the factors influencing the erosion, the path analysis was performed to differentiate direct and indirect effects of water and vegetation factors on the processes of soil erosion by water (Table S5).

**Table S4.** Correlation matrix between soil erosion processes (i.e., changing area of soil erosion) and seven water and vegetation factors during the period of 1978-2008.

| Variable | Soil erosion processes | Changes in amount of conifer | Changes in amount of broad-leaved | Changes in amount of mixed forests | Changes in amount of shrubs | Changes in amount of grass | Precipitation | Changes in amount of evapotranspiration |
| --- | --- | --- | --- | --- | --- | --- | --- | --- |

|  |  | forests | forests |  |  |  |  |  |
| --- | --- | --- | --- | --- | --- | --- | --- | --- |
| <u>During the period of 1978-1990</u> |  |  |  |  |  |  |  |  |
| Soil erosion processes | 1.00 |  |  |  |  |  |  |  |
| Changes in amount of coniferous forests | 0.17* | 1.00 |  |  |  |  |  |  |
| Changes in amount of broadleaved forests | 0.41** | 0.20* | 1.00 |  |  |  |  |  |
| Changes in amount of mixed forests | 0.09 | -0.06 | -0.13 | 1.00 |  |  |  |  |
| Changes in amount of shrubs | 0.15* | 0.00 | -0.03 | 0.01 | 1.00 |  |  |  |
| Changes in amount of grass | -0.11* | 0.02 | -0.07 | -0.01 | -0.14 | 1.00 |  |  |
| Precipitation | 0.10 | -0.01 | -0.02 | 0.14* | 0.09 | 0.52** | 1.00 |  |
| Changes in amount of evapotranspiration | 0.12* | -0.05 | -0.03 | 0.08 | -0.06 | 0.37 | 0.71** | 1.00 |
| <u>During the period of 1990-2000</u> |  |  |  |  |  |  |  |  |
| Soil erosion processes | 1.00 |  |  |  |  |  |  |  |
| Changes in amount of coniferous forests | 0.11* | 1.00 |  |  |  |  |  |  |
| Changes in amount of broadleaved forests | 0.34** | 0.04 | 1.00 |  |  |  |  |  |
| Changes in amount of mixed forests | 0.07 | 0.00 | 0.11 | 1.00 |  |  |  |  |
| Changes in amount of shrubs | -0.12* | 0.04 | 0.00 | 0.00 | 1.00 |  |  |  |
| Changes in amount of grass | 0.09 | 0.00 | -0.12* | 0.01 | -0.21 | 1.00 |  |  |
| Precipitation | 0.18* | 0.05 | 0.00 | 0.05 | 0.12 | -0.23* | 1.00 |  |
| Changes in amount of evapotranspiration | 0.22** | -0.08 | -0.16* | -0.02 | -0.51** | -0.32** | 0.47** | 1.00 |
| <u>During the period of 2000-2008</u> |  |  |  |  |  |  |  |  |
| Soil erosion processes | 1.00 |  |  |  |  |  |  |  |
| Changes in amount of coniferous forests | 0.12* | 1.00 |  |  |  |  |  |  |
| Changes in amount of broadleaved forests | 0.38** | 0 | 1.00 |  |  |  |  |  |
| Changes in amount of mixed forests | -0.09 | 0 | 0.02 | 1.00 |  |  |  |  |
| Changes in amount of shrubs | 0.15* | 0.02 | 0 | 0 | 1.00 |  |  |  |
| Changes in amount of grass | -0.10* | -0.02 | -0.01 | 0 | -0.22* | 1.00 |  |  |
| Precipitation | -0.19* | 0.05 | 0.03 | -0.08 | 0.14* | 0.32** | 1.00 |  |
| Changes in amount of evapotranspiration | 0.34** | -0.08 | -0.05 | -0.04 | -0.07 | 0.35** | 0.61** | 1.00 |
| <u>During the period of 1978-2008</u> |  |  |  |  |  |  |  |  |
| Soil erosion processes | 1.00 |  |  |  |  |  |  |  |
| Changes in amount of coniferous forests | 0.21* | 1.00 |  |  |  |  |  |  |
| Changes in amount of broadleaved forests | 0.40** | -0.1 | 1.00 |  |  |  |  |  |
| Changes in amount of mixed forests | 0.09 | 0.01 | 0.02 | 1.00 |  |  |  |  |
| Changes in amount of shrubs | 0.15* | -0.03 | 0 | 0 | 1.00 |  |  |  |
| Changes in amount of grass | 0.08 | 0.04 | -0.09 | -0.01 | -0.16* | 1.00 |  |  |
| Precipitation | -0.11* | 0.03 | -0.11* | 0 | 0.12* | 0.57** | 1.00 |  |
| Changes in amount of evapotranspiration | 0.19* | -0.08 | -0.07 | -0.01 | 0.09 | 0.62** | 0.76** | 1.00 |

\* Significant at $P<0.05$ , \*\* Significant at $P<0.01$ .

**Table S5.**
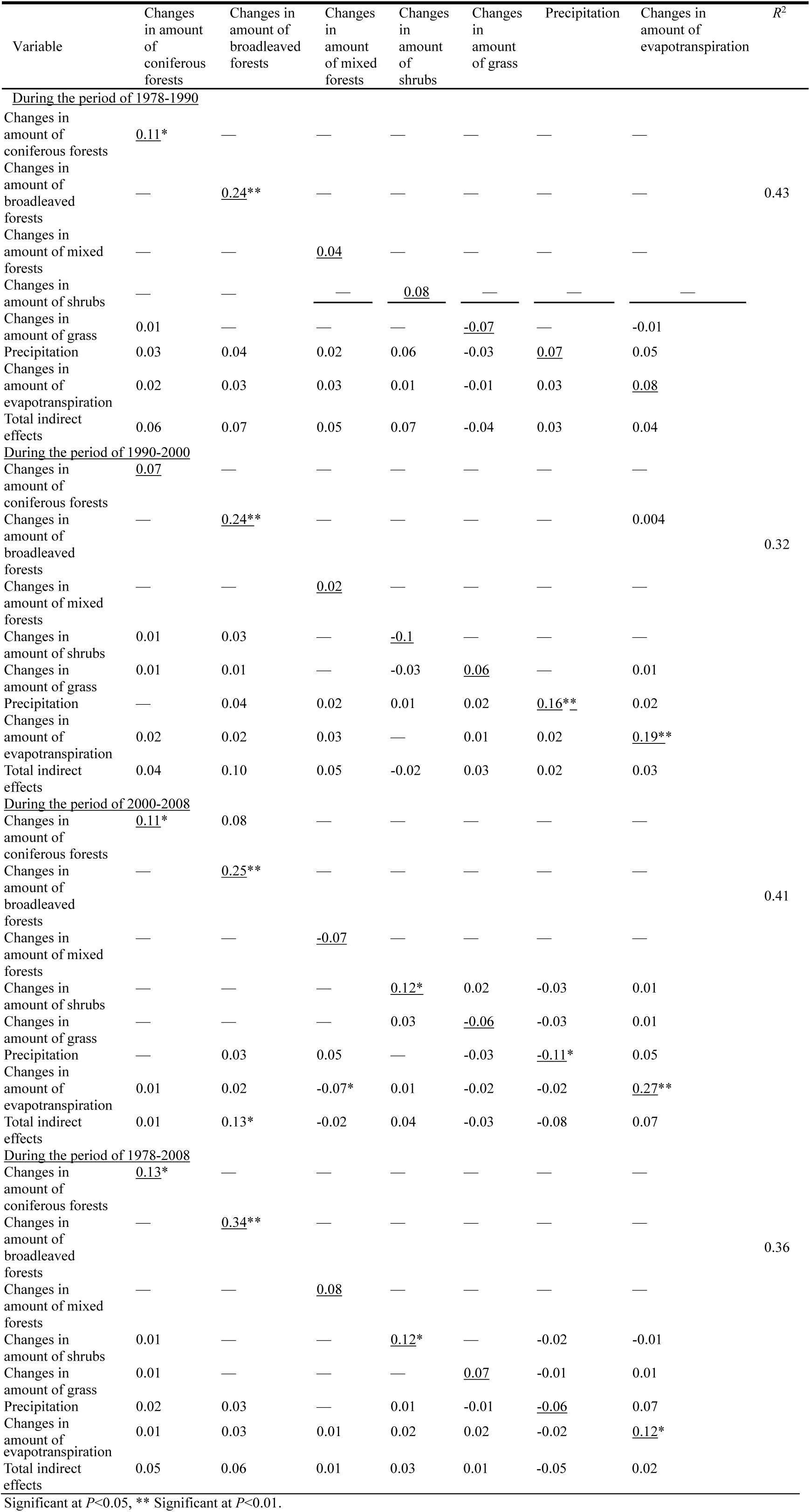
Direct effects (diagonal, underlined), indirect effects (off-diagonal), and total indirect effects of factors on the soil erosion processes (n=860) during the period of 1978-2008.

### Impact on desertification

#### 1) Monitoring of desertification

Desertification intensities were defined as extremely severe, severe, moderate, and slight (Table S6) (47). A series of Landsat MSS, TM and ETM+ images taken between 1978 and 2008 were obtained. As it was difficult to acquire cloud-free images that covered the whole study area within a given year because of the large geographical range covered by the study region, some images from previous or subsequent years were selected to replace unsuitable images from 1978 (1977-1980), 1990’s, 2000’s and 2008 (2007-2009). The spatial resolutions of the TM/ETM+ and MSS data are 30 and 80 m, respectively. All images represent composites of bands 4, 3, and 2 (R, G, and B, respectively) and were used to create false-color images.

Visual interpretation was used to estimate the area of the aeolian desertification by each intensity level. Our estimation achieved an overall accuracy of more than 90.0% based on field survey (3100 plots with 30 m×30 m).

**Table S6.**
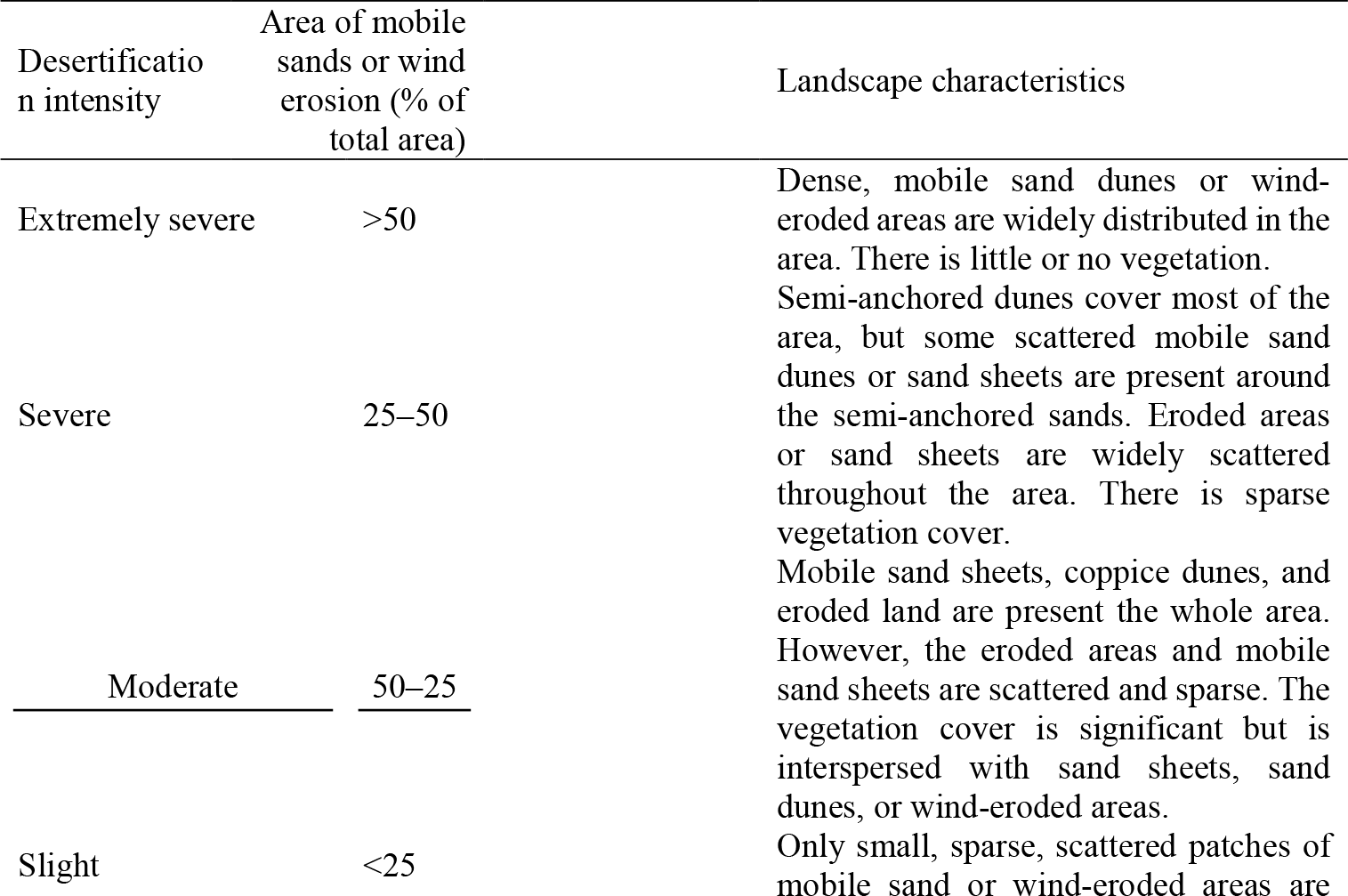
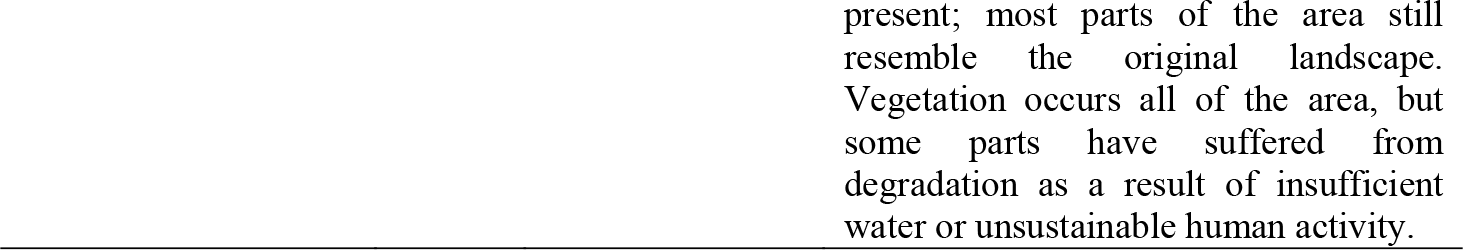
Indices of desertification used for the classification of desertified land.

| Desertification intensity | Area of mobile sands or wind erosion (% of total area) | Landscape characteristics |
| --- | --- | --- |
| Extremely severe | >50 | Dense, mobile sand dunes or wind-eroded areas are widely distributed in the area. There is little or no vegetation. |
| Severe | 25–50 | Semi-anchored dunes cover most of the area, but some scattered mobile sand dunes or sand sheets are present around the semi-anchored sands. Eroded areas or sand sheets are widely scattered throughout the area. There is sparse vegetation cover. |
| Moderate | 50–25 | Mobile sand sheets, coppice dunes, and eroded land are present the whole area. However, the eroded areas and mobile sand sheets are scattered and sparse. The vegetation cover is significant but is interspersed with sand sheets, sand dunes, or wind-eroded areas. |
| Slight | <25 | Only small, sparse, scattered patches of mobile sand or wind-eroded areas are |

#### 2) The spatial distribution of forests and shrublands in desertification regions

The spatial distribution of desertification in 1978 was the initial state of desertification in the TNAP. Due to the change of desertification was caused by so many factors, we defined the original spatial distribution of desertification, i.e., the spatial distribution of desertification in 1978, as the reference to illustrate the contribution rate of forests and shrublands. We obtained the spatial distribution of desertification in 1978 based on Landsat MSS remote sensing data. Then, we obtained the spatial distributions of forests and shrublands in desertification regions by overlaying the maps of forests and shrublands from 1978 to 2008 on the map of desertification regions in 1978.

#### 3) Estimation of the impact on desertification

The contribution rates of forests and shrublands to decrease of desertification area were defined as the reduction of desertification area by considering only the direct effect from site occupancy by forests and shrublands. Because the indirect effect (the shelter impacts of forests and shrublands) was not considered, our estimation on the contribution of forests and shrublands was somewhat conservative.

**Table AS1.** Summary of the investment from the government and labor cost. The 2016 values were derived from CPI inflation calculator from US Department of Labor, Bureau of Labor Statistics (http://www.bls.gov/data/inflation_calculator.htm; accessed 12/08/2016).

| Year | Investment<br>(Million<br>RMB) | Investment<br>(Million<br>USD) | Investment<br>in 2016<br>value<br>(Million<br>USD) | Labor<br>cost<br>(Million<br>RMB)* | Labor<br>cost<br>(Million<br>USD)* | Labor cost<br>in 2016 value<br>(Million<br>USD)* | Total cost<br>(Million<br>RMB)** | Total<br>cost<br>(Million<br>USD)** | Total cost in<br>2016 value<br>(Million<br>USD)** | Exchange<br>rate of<br>USD to<br>RMB |
| --- | --- | --- | --- | --- | --- | --- | --- | --- | --- | --- |
| 1979 | 44.77 | 28.89 | 96.19 | 9133.63<br>(actual<br>labor cost<br>from 1979<br>to 1985) | 4228.53<br>(mean<br>exchange<br>rates:<br>1:2.16) | 11131.81<br>(mean<br>exchange<br>rates: 1:2.16) | 1349.57 | 632.97 | 2107.54 | 1:1.55 |
| 1980 | 50.99 | 34.22 | 100.39 |  |  |  | 1355.79 | 638.30 | 1872.52 | 1:1.49 |
| 1981 | 45.22 | 25.45 | 67.68 |  |  |  | 1350.02 | 629.53 | 1674.10 | 1:1.78 |
| 1982 | 53.35 | 27.72 | 69.44 |  |  |  | 1358.15 | 631.80 | 1582.64 | 1:1.92 |
| 1983 | 48.75 | 24.91 | 60.46 |  |  |  | 1353.55 | 628.99 | 1526.65 | 1:1.96 |
| 1984 | 62.87 | 28.52 | 66.35 |  |  |  | 1367.67 | 632.60 | 1471.77 | 1:2.20 |
| 1985 | 60.29 | 20.53 | 46.12 |  |  |  | 1365.09 | 624.61 | 1403.22 | 1:2.94 |
| 1986 | 49.09 | 14.22 | 31.35 | 19599.78<br>(total<br>investmen<br>t of labor<br>from 1986<br>to 1995) | 3698.07<br>(mean<br>exchange<br>rates:<br>1:5.30) | 6864.98<br>(mean<br>exchange<br>rates: 1:5.30) | 2009.07 | 384.03 | 846.99 | 1:3.45 |
| 1987 | 52.19 | 14.02 | 29.83 |  |  |  | 2012.17 | 383.83 | 816.73 | 1:3.72 |
| 1988 | 55.73 | 14.97 | 30.59 |  |  |  | 2015.71 | 384.78 | 786.24 | 1:3.72 |
| 1989 | 53.67 | 14.25 | 27.78 |  |  |  | 2013.65 | 384.06 | 748.70 | 1:3.77 |
| 1990 | 167.33 | 34.98 | 64.7 |  |  |  | 2127.31 | 404.79 | 748.66 | 1:4.78 |
| 1991 | 197.50 | 37.10 | 65.85 |  |  |  | 2157.48 | 406.91 | 722.19 | 1:5.32 |
| 1992 | 249.21 | 45.19 | 77.86 |  |  |  | 2209.19 | 415.00 | 715.02 | 1:5.51 |
| 1993 | 350.80 | 60.88 | 101.86 |  |  |  | 2310.78 | 430.69 | 720.50 | 1:5.76 |
| 1994 | 389.28 | 45.17 | 73.68 |  |  |  | 2349.26 | 414.98 | 676.88 | 1:8.62 |
| 1995 | 424.59 | 50.84 | 80.64 |  |  |  | 2384.57 | 420.65 | 667.21 | 1:8.35 |
| 1996 | 711.69 | 85.60 | 131.88 | 1010.28 | 121.57 | 187.30 | 1721.97 | 207.17 | 319.18 | 1:8.31 |
| 1997 | 805.67 | 97.19 | 146.38 | 1029.35 | 124.17 | 187.10 | 1835.02 | 221.36 | 333.48 | 1:8.29 |
| 1998 | 902.89 | 109.06 | 161.74 | 1177.92 | 142.26 | 210.97 | 2080.81 | 251.32 | 372.71 | 1:8.28 |
| 1999 | 1187.54 | 143.45 | 208.14 | 1125.79 | 135.96 | 197.27 | 2313.33 | 279.41 | 405.41 | 1:8.28 |
| 2000 | 1303.46 | 157.45 | 221.02 | 1423.11 | 171.87 | 241.27 | 2726.57 | 329.32 | 462.29 | 1:8.28 |
| 2001 | 918.72 | 111.00 | 151.51 | 792.16 | 95.67 | 130.58 | 1710.88 | 206.67 | 282.09 | 1:8.28 |
| 2002 | 932.07 | 112.61 | 151.31 | 574.64 | 69.40 | 93.25 | 1506.71 | 182.01 | 244.56 | 1:8.28 |
| 2003 | 849.07 | 102.58 | 134.76 | 356.71 | 43.08 | 56.60 | 1205.78 | 145.66 | 191.36 | 1:8.28 |
| 2004 | 825.10 | 99.69 | 127.57 | 358.04 | 43.24 | 55.33 | 1183.14 | 142.93 | 182.90 | 1:8.28 |
| 2005 | 896.35 | 109.42 | 135.43 | 384.06 | 46.89 | 58.04 | 1280.41 | 156.31 | 193.47 | 1:8.19 |
| 2006 | 816.18 | 102.38 | 122.76 | 330.92 | 41.52 | 49.78 | 1147.1 | 143.90 | 172.54 | 1:7.97 |
| 2007 | 797.56 | 104.94 | 122.34 | 317.09 | 41.72 | 48.64 | 1114.65 | 146.66 | 170.98 | 1:7.60 |
| 2008 | 1946.25 | 280.23 | 314.63 | 364.59 | 52.46 | 58.90 | 2310.84 | 332.69 | 373.53 | 1:6.95 |
| **Sum** | **15248.18** | **2137.46** | **3220.24** | **37978.07** | **9056.44** | **19571.82** | **53226.24** | **11193.93** | **22792.06** |  |

**Table AS2.** Examples of tree species used for afforestation were poorly matched to local site conditions in the TNAP region

| Location | Precipitation<br>(mm) | Species | Planting density<br>(trees ha<sup>-1</sup>) |
| --- | --- | --- | --- |
| Zhangbei County,<br>40°57'~41°34'N,<br>114°10'~115°27'E | 309~480 | *Populus* spp. | 1667 |
| Tongliao City,<br>42°15'~45°41'N,<br>119°15'~123°43'E | 343~500 | *Populus* spp. | 2498 |
| Naiman County,<br>42°14'~43°32'N,<br>120°19'~121°35'E | 305~475 | *Pinus*<br>*sylvestris* var. *mongolica* | 2300~4400 |
| Zhangwu County,<br>42°07'~42°51'N,<br>121°53'~122°58'E | 474 | *Pinus*<br>*sylvestris* var. *mongolica* | 3300~6600 |
| West of Liaoning<br>Province,<br>40°20'~42°34'N,<br>118°50'~120°15'E | 400~600 | *Pinus tabulaeformis* | 3300~4400 |
| Wutai County,<br>38°50'~39°05'N,<br>113°29'~113°44'E | 400~500 | *Larix principis-rupprechtii* | 2500~4400 |
| Hexi Corridor,<br>37°57'~39°47'N,<br>101°09'~103°54'E | 115~390 | *Populus* spp. | 3300 |
| Dehui County,<br>44°02'~44°53'N,<br>125°14'~126°24'E | 520 | *Populus* spp. | 1667~3300 |

**Table AS3.**
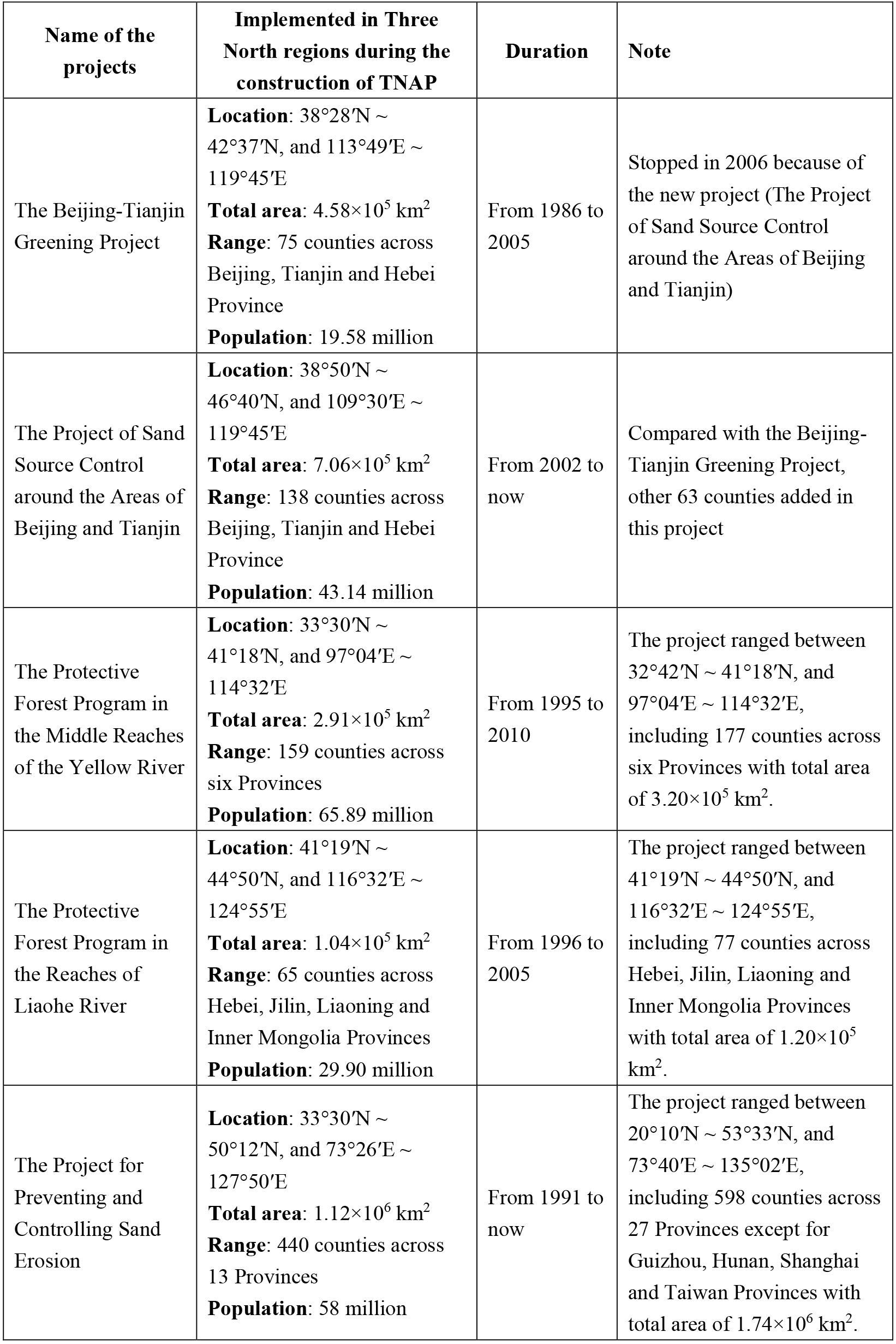
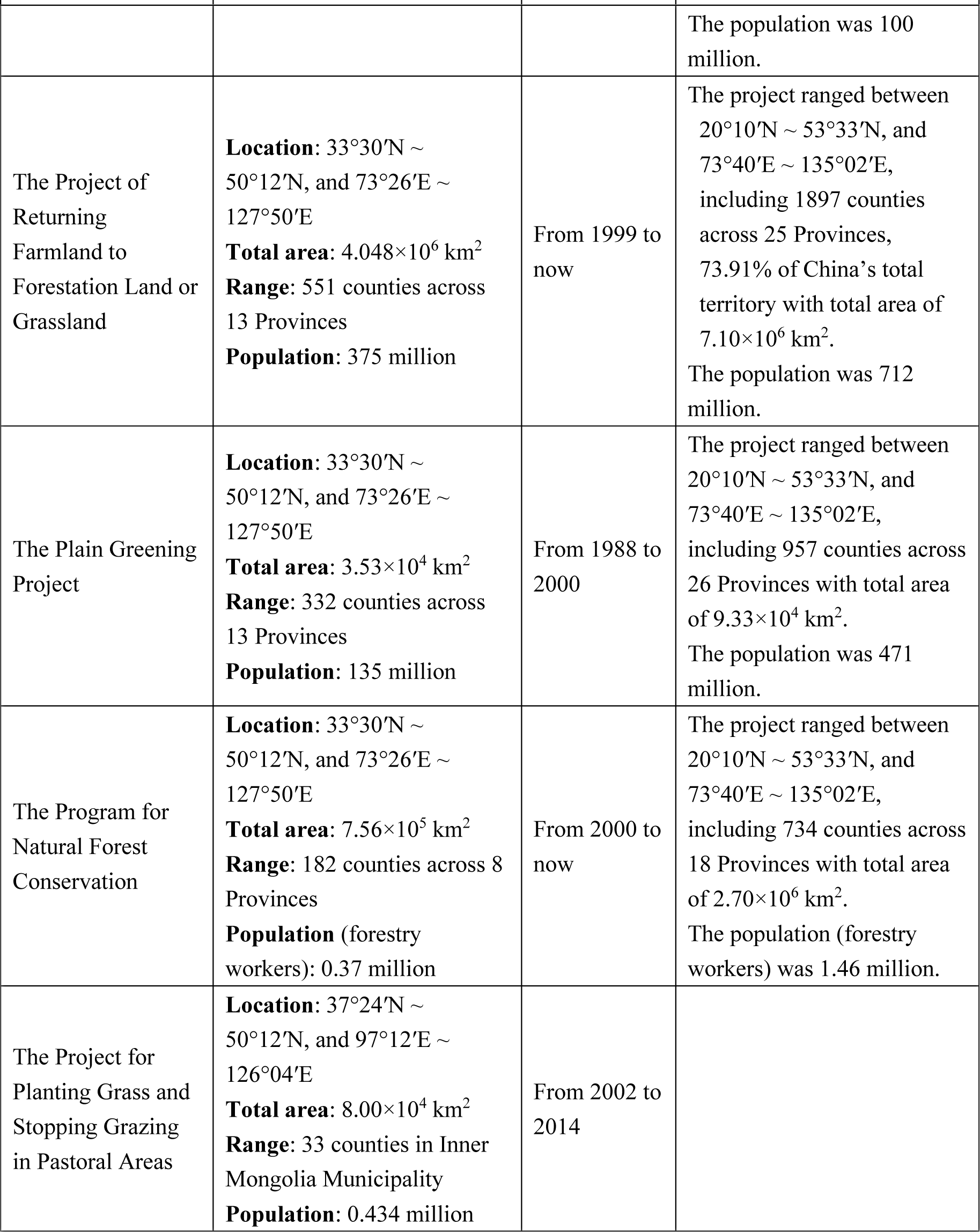
List of the projects implemented in the Three North regions during theconstruction of TNAP

